# Motivated semantic control: Exploring the effects of extrinsic reward and self-reference on semantic retrieval in semantic aphasia

**DOI:** 10.1101/2021.05.25.444996

**Authors:** Nicholas E. Souter, Sara Stampacchia, Glyn Hallam, Hannah Thompson, Jonathan Smallwood, Elizabeth Jefferies

## Abstract

Recent insights show increased motivation can benefit executive control, but this effect has not been explored in relation to semantic cognition. Patients with deficits of controlled semantic retrieval in the context of semantic aphasia (SA) after stroke may benefit from this approach since ‘semantic control’ is considered an executive process. Deficits in this domain are partially distinct from domain-general deficits of cognitive control. We assessed the effect of both extrinsic and intrinsic motivation in healthy controls and semantic aphasia patients. Experiment 1 manipulated extrinsic reward using high or low levels of points for correct responses during a semantic association task. Experiment 2 manipulated the intrinsic value of items using self-reference; allocating pictures of items to the participant (‘self’) or researcher (‘other’) in a shopping game before people retrieved their semantic associations. These experiments revealed that patients, but not controls, showed better performance when given an extrinsic reward, consistent with the view that increased external motivation may help to ameliorate patients’ semantic control deficits. However, while self-reference was associated with better episodic memory, there was no effect on semantic retrieval. We conclude that semantic control deficits can be reduced when extrinsic rewards are anticipated; this enhanced motivational state is expected to support proactive control, for example, through the maintenance of task representations. It may be possible to harness this modulatory impact of reward to combat the control demands of semantic tasks in SA patients.

## 1. Introduction

Our ability to understand the world relies on flexible access to conceptual information within a semantic store (Jefferies, 2013). Evidence supports the existence of dissociable systems underlying the storage and retrieval of semantic representations (Lambon Ralph et al., 2017). Semantic dementia patients with relatively focal atrophy focussed on the ventrolateral anterior temporal lobes show degraded semantic knowledge, while patients with semantic aphasia (SA) experience deregulated semantic retrieval, or semantic control, following left prefrontal and/or temporoparietal stroke (Jefferies & Lambon Ralph, 2006). Semantic control is an executive process which supports the retrieval of non-dominant aspects of knowledge whilst overcoming competition from distractors (Hoffman et al., 2018; Jefferies, 2013). Impaired semantic control in SA gives rise to deficits in both verbal communication and organisation of nonverbal actions (Jefferies et al., 2019), consistent with the definition of SA as impaired manipulation of verbal and non-verbal symbolic information (Head, 1926). In line with the damage to left ventrolateral prefrontal cortex and/or left temporoparietal regions in SA, studies of healthy participants employing neuroimaging (Jackson, 2021) and transcranial magnetic stimulation (Hallam et al., 2016) have implicated both left inferior frontal gyrus (LIFG) and posterior middle temporal gyrus (pMTG) in semantic control.

In SA, access to semantic knowledge is not universally compromised, but depends on task demands. Semantic retrieval is impaired for subordinate meanings, and when inhibition of task-irrelevant distractors is required (Jefferies, 2013). This results in reduced flexibility when retrieving semantic information in ambiguous contexts (Noonan et al., 2010). Impaired semantic control in SA is also evident when retrieving thematic associations between concepts: identifying weak as opposed to strong associations requires semantic control processes that focus retrieval on non-dominant conceptual information (Thompson et al., 2017). Research has explored manipulations which ameliorate semantic control deficits in SA, such as cueing. Successive phonemic cues (e.g. c.. ca.. cam.. for CAMEL) can facilitate picture naming (Jefferies et al., 2008), while contextually-relevant sentences (Noonan et al., 2010) or emotional and location cues (Lanzoni et al., 2019) can facilitate the retrieval of non-dominant interpretations of ambiguous homonyms. Cues reduce control demands by narrowing down the number of retrievable options and biasing retrieval towards task-relevant information.

An alternative approach to facilitating semantic retrieval involves recruiting processes beyond semantic cognition. Investigations with healthy adults have demonstrated that extrinsic rewards, such as monetary incentives or awarded points, can improve performance in domains including control of visual attention (Padmala & Pessoa, 2011), task-switching (Capa et al., 2013), contextual control (Kouneiher et al., 2009), creative problem solving (Cristofori et al., 2018), interference control (Zhao et al., 2018), and conflict adaptation (Dreisbach & Fischer, 2012). Behavioural benefits of extrinsic reward include increased accuracy, reduced reaction times, and reduced switch-costs (Yee & Braver, 2018). Extrinsic incentives are considered a key element of ‘gamification’ (Mekler et al., 2017), which uses typical elements of digital games to increase engagement with training activities including post-stroke rehabilitation (Romani et al., 2019). To our knowledge, extrinsic rewards have not been used previously in conjunction with semantic tasks or in SA patients.

Tasks with high control demands are effortful as they draw on limited resources including selective attention and working memory (Yee & Braver, 2018). The cost of mental exertion is typically weighed against the potential benefits of the action (Botvinick & Braver, 2015). As such, tasks perceived as high effort and low in reward may be less appealing than more trivial low effort and high reward actions. Introducing task-based incentives can offset perceived costs (Goschke & Bolte, 2014) and increase preparatory control, and therefore one’s ability to sustainably engage with a task (Notebaert & Braem, 2015). This can benefit either cognitive stability or flexibility, depending on recent reward history (Fröber et al., 2019). The neural processing of extrinsic reward has been consistently linked to a network of regions including the ventromedial prefrontal cortex, caudate, and putamen (Lin et al., 2012). Cumulative reward value appears to be tracked and represented in these regions (Juechems et al., 2017). Effects of reward on cognition have been attributed to dopaminergic transmission between these regions and the multiple demand network (MDN), which supports challenging tasks across domains (Camilleri et al., 2018; Parro et al., 2018).

While extrinsic reward refers to incentives provided externally, intrinsic reward refers to inherent enjoyment of or interest in a task (Mori et al., 2018). Intrinsic motivation is relatively difficult to manipulate experimentally, but can be modulated indirectly, through factors such as self-reference. Tamir and Mitchell (2012) demonstrated that self-referential information is intrinsically motivating; participants reliably choose to forgo monetary incentives in order to disclose information about the self, in conjunction with increased activation in brain regions associated with reward processing. The neural substrates underlying both intrinsic motivation and self-reference show considerable overlap with reward circuitry (Di Domenico & Ryan, 2017; Enzi et al., 2009). Cognition shows biases in favour of self-referenced items, within perception (Sui et al., 2012), attention (Sui & Humphreys, 2015b), working memory (Röer et al., 2013), and recognition memory (Hou et al., 2019). Moreover, Sui and Humphreys (2015a) demonstrated that extrinsic reward and self-reference confer separable but equivalent benefits in associative learning. Self-reference benefits to episodic memory persist in patients with SA (Stampacchia et al., 2019).

If regions associated with reward processing are intact it may be possible to harness modulatory effects of motivation in rehabilitation for post-stroke aphasia. The benefits of increased motivation may be more pronounced in more impaired patients, with greater difficulties constraining internal representations increasing reliance on external prompts. Given evidence of effects of reward and self-reference on cognitive functions across domains, similar benefits may occur for semantic control. However, semantic control is dissociable from domain-general control: the peak activations in fMRI studies that manipulate semantic control demands in healthy participants fall outside the multiple-demand network (Gao et al., 2021; Noonan et al., 2013), and inhibitory stimulation of semantic control sites temporarily disrupts control-demanding semantic tasks but not demanding visual judgements (Whitney et al., 2011). Moreover, while impaired semantic control is ubiquitous in semantic aphasia, some but not all of these patients have general deficits of cognitive control: lesion-symptom mapping shows that these semantic and non-semantic control deficits are associated with different patterns of structural damage (Souter, Wang, et al., 2021). Given this distinction, there is a need to investigate the effects of motivation in tasks with high semantic control demands, to establish if this domain can benefit from ‘gamification’ strategies to the same degree as other cognitive tasks. Furthermore, evidence suggests affective abnormalities in SA, including in the ability to categorise facial portrayals according to discrete emotion categories (Souter, Lindquist, & Jefferies, 2021). This is thought to reflect deficits in constraining internal states beyond the conceptual domain, which may extend to, and therefore limit modulatory effects of, motivation. While people with aphasia generally benefit from the use of motivating ‘gamification’ strategies, we cannot assume that these benefits will transfer to people with SA for this reason. This is a key motivation for the current study. SA patients have been shown to benefit from the provision of external cues that provide additional information pertaining to semantic decisions (Noonan et al., 2010), but the influence of reward manipulations, which provide external prompts in the absence of contextually-relevant information, have not been investigated to our knowledge.

The current study aimed to investigate the influence of both extrinsic reward and intrinsic motivation induced through self-reference on the retrieval of strong and weak thematic associations in SA. Experiment 1 assessed the effect of cued extrinsic reward, in the form of high or low levels of performance-contingent token points. Experiment 2 assessed the effect of self-reference by allocating pictured ‘shopping items’ either to the participant (‘self’ condition) or the researcher (‘other’ condition), prior to semantic judgements about these items. If modulatory effects of motivation can benefit semantic control, high extrinsic reward and/or self-reference might ameliorate patients’ semantic deficits. Motivation may support the maintenance of task goals when semantic control is deficient; consequently, any performance gains would likely be greater for weak associations, which place higher demands on semantic control. A better understanding of the effects of motivation on semantic retrieval of strong and weak associations may have implications for the use of gamified approaches for aphasia rehabilitation.

## 2. Method

### 2.1. Participants

The sample consisted of 16 SA patients (nine females), and 15 controls (12 females). All participated in Experiment 1, while a subset participated in Experiment 2 (demographic information is presented separately in *3.1.1.* and *4.1.1.*, respectively). Patients were recruited from communication support groups across *ANONYMISED*. All had aphasia following left hemisphere stroke and were at least 18 months post-stroke. Patients were selected to show impairments in both verbal and non-verbal semantic cognition, consistent with previous definitions of SA (Jefferies & Lambon Ralph, 2006). The criteria used are explained in the supplementary section *Background Neuropsychology*. Controls were healthy adults matched to the patients on age and years in education and reported no history of psychiatric or neurological disorders. Informed consent was obtained for all participants.

### 2.2. Lesion analyses

Ten patients (P1 – P10) had MRI scans at *ANONYMISED* (see supplementary section *MRI Acquisition* for scanning protocols and extraction/registration procedure). All ten participated in Experiment 1, four (P3 – P6) participated in Experiment 2. Each patient’s lesion was manually traced in MRIcron. Figure 1a provides a lesion overlap map for these patients. Eight cases showed damage to LIFG. Several patients showed damage to other regions including the pMTG, superior temporal gyrus, and supramarginal gyrus. Clinical acute-stage scans were available for two further patients and revealed damage to LIFG (MRI for P16) and a left frontoparietal lesion (CT for P12). Lesion information was not available for the remaining four patients due to contraindications for scanning and/or closure of scanning facilities during the COVID-19 pandemic.

**Figure 1.**
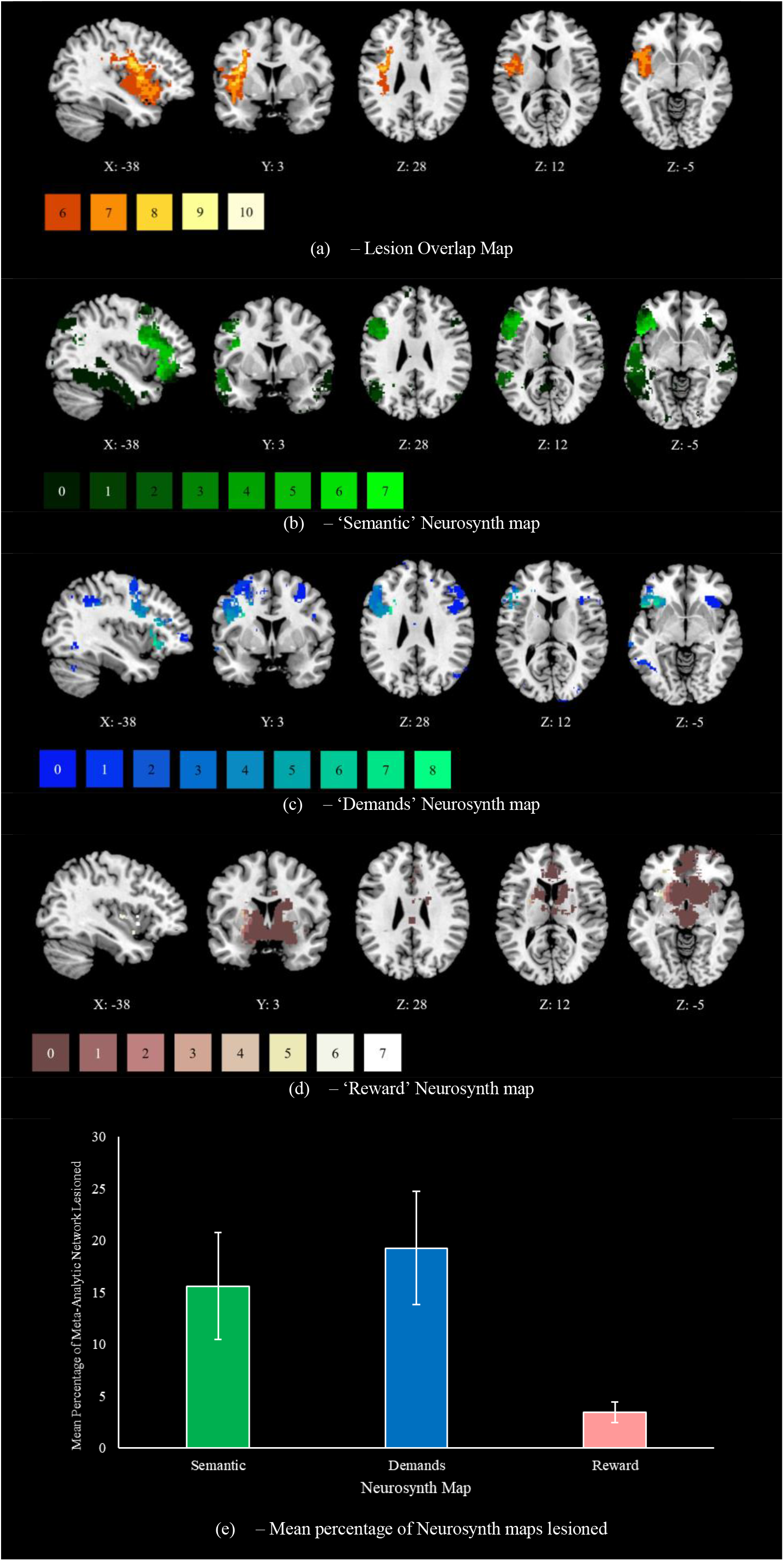
Patient lesion analyses, including (a) a lesion overlap map for ten semantic aphasia patients in the current study, created using manual segmentation in MRICron. This map shows lesion overlap in 6 or more patients, with the colour of the lesioned area corresponding to the number of affected cases (bottom left). We assessed the extent of overlap between patient lesions and term-based meta-analytic maps from Neurosynth for the terms (b) ‘semantic’ (1031 studies), (c) ‘demands’ (596 studies), and (d) ‘reward’ (922 studies). Neurosynth maps are coloured according to impact by lesion across the sample, with brighter areas reflecting those more often implicated in lesions. Each map was formatted in MRICron. We present (e) the mean percentage of each map lesioned across patients, with standard error of the mean error bars.

To assess the impact of patients’ lesions on functional networks of interest, we extracted maps from Neurosynth using term-based meta-analyses (Yarkoni et al., 2011) for ‘Semantic’, ‘Demands’, and ‘Reward’ (Figure 1b-d). This allowed us to observe the extent to which patients present with damage to regions associated with semantic processing, domain-general task demands, and reward processing. We calculated the average percentage of each map that was damaged across the patients with available lesion maps. Analysis was restricted to left hemisphere aspects of each network, such that it reflects a percentage of all voxels that could possibly be lesioned in an exclusively left-hemisphere stroke sample. While right-hemisphere aspects of these networks may be affected by disconnection (Souter, Wang, et al., 2021), this is beyond the scope of the current paper. As seen in Figure 1e, patients showed the most damage to ‘Demands’ regions, followed by ‘Semantic’ regions. ‘Reward’ regions were relatively spared, suggesting it might be possible to harness modulatory impacts of reward.

### 2.3. Background neuropsychological testing

Patients completed background tests of language, memory, executive function, and semantic cognition. The control participants tested on the experimental tasks in this study did not complete these background assessments. Patients’ individual scores are provided in Table 1 for background neuropsychology and Table 2 for semantic tests. Interpretation of the sample’s performance is provided in the supplementary *Background Neuropsychology* section. Patients presented with variable levels of impairment in speech fluency and word repetition. Most patients presented with impaired working memory. Visuospatial processing was largely preserved. Eleven patients showed impairment on at least one test of executive function.

**Table 1.**
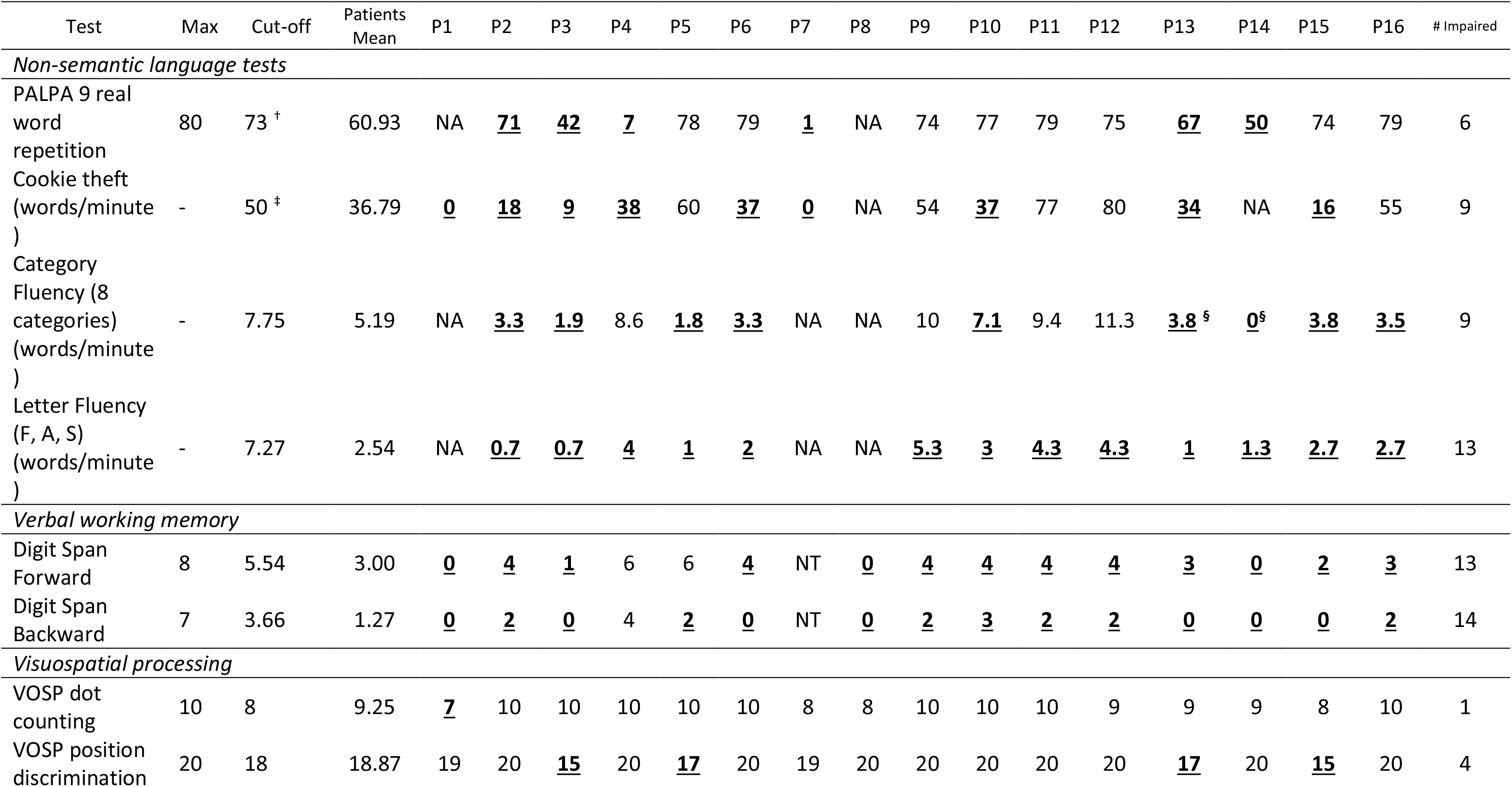
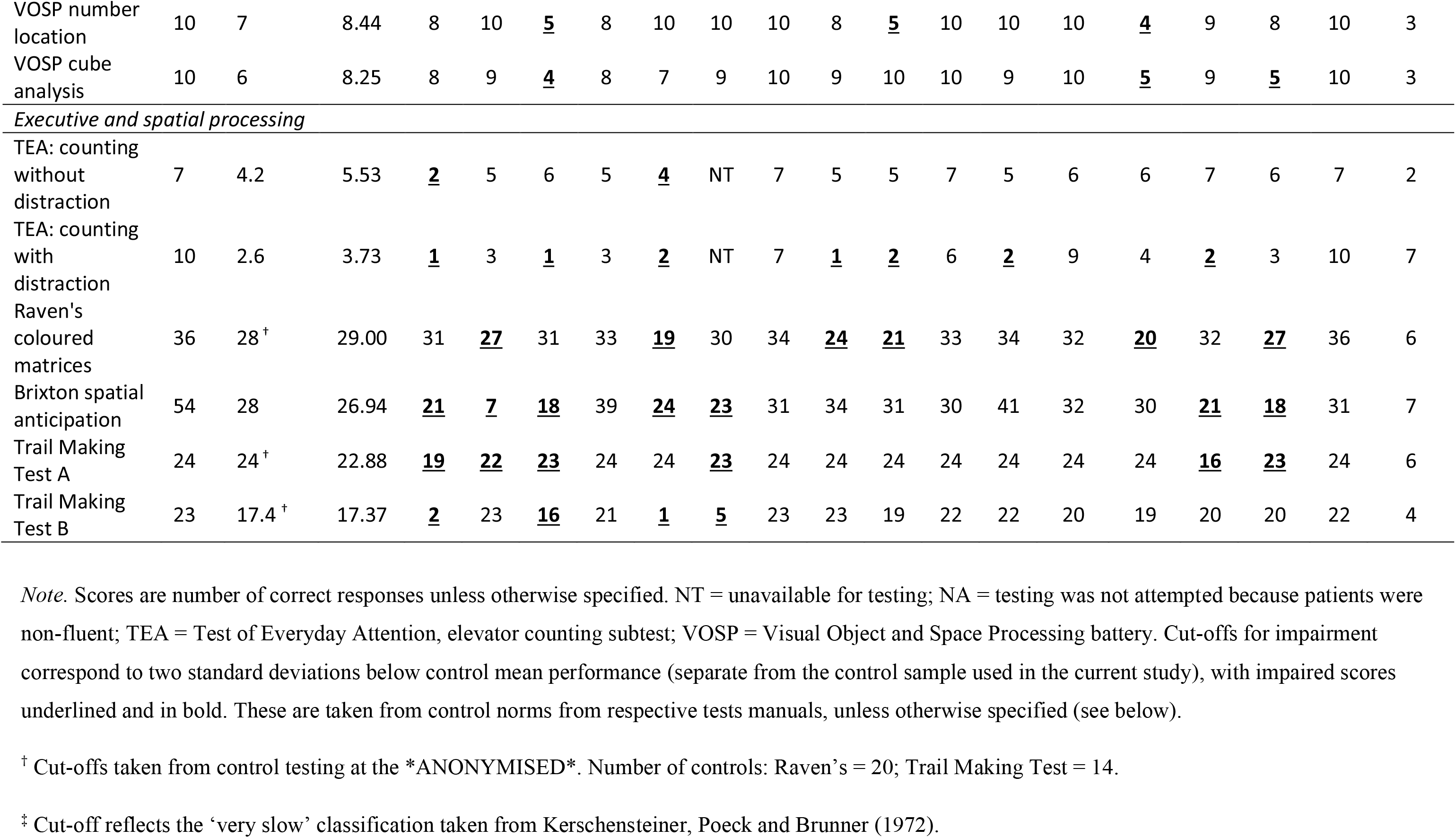
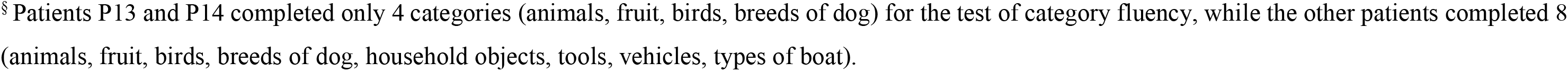
Patient performance on background neuropsychological testing.

| Test | Max | Cut-off | Patients Mean | P1 | P2 | P3 | P4 | P5 | P6 | P7 | P8 | P9 | P10 | P11 | P12 | P13 | P14 | P15 | P16 | # Impaired |
| --- | --- | --- | --- | --- | --- | --- | --- | --- | --- | --- | --- | --- | --- | --- | --- | --- | --- | --- | --- | --- |
| *Non-semantic language tests* |  |  |  |  |  |  |  |  |  |  |  |  |  |  |  |  |  |  |  |  |
| PALPA 9 real word repetition | 80 | 73 <sup>†</sup> | 60.93 | NA | <u>71</u> | <u>42</u> | <u>7</u> | 78 | 79 | <u>1</u> | NA | 74 | 77 | 79 | 75 | <u>67</u> | <u>50</u> | 74 | 79 | 6 |
| Cookie theft (words/minute ) | - | 50 <sup>‡</sup> | 36.79 | <u>0</u> | <u>18</u> | <u>9</u> | <u>38</u> | 60 | <u>37</u> | <u>0</u> | NA | 54 | <u>37</u> | 77 | 80 | <u>34</u> | NA | <u>16</u> | 55 | 9 |
| Category Fluency (8 categories) (words/minute ) | - | 7.75 | 5.19 | NA | <u>3.3</u> | <u>1.9</u> | 8.6 | <u>1.8</u> | <u>3.3</u> | NA | NA | 10 | <u>7.1</u> | 9.4 | 11.3 | <u>3.8</u> <sup>§</sup> | <u>0</u> <sup>§</sup> | <u>3.8</u> | <u>3.5</u> | 9 |
| Letter Fluency (F, A, S) (words/minute ) | - | 7.27 | 2.54 | NA | <u>0.7</u> | <u>0.7</u> | <u>4</u> | <u>1</u> | <u>2</u> | NA | NA | <u>5.3</u> | <u>3</u> | <u>4.3</u> | <u>4.3</u> | <u>1</u> | <u>1.3</u> | <u>2.7</u> | <u>2.7</u> | 13 |
| *Verbal working memory* |  |  |  |  |  |  |  |  |  |  |  |  |  |  |  |  |  |  |  |  |
| Digit Span Forward | 8 | 5.54 | 3.00 | <u>0</u> | <u>4</u> | <u>1</u> | 6 | 6 | <u>4</u> | NT | <u>0</u> | <u>4</u> | <u>4</u> | <u>4</u> | <u>4</u> | <u>3</u> | <u>0</u> | <u>2</u> | <u>3</u> | 13 |
| Digit Span Backward | 7 | 3.66 | 1.27 | <u>0</u> | <u>2</u> | <u>0</u> | 4 | <u>2</u> | <u>0</u> | NT | <u>0</u> | <u>2</u> | <u>3</u> | <u>2</u> | <u>2</u> | <u>0</u> | <u>0</u> | <u>0</u> | <u>2</u> | 14 |
| *Visuospatial processing* |  |  |  |  |  |  |  |  |  |  |  |  |  |  |  |  |  |  |  |  |
| VOSP dot counting | 10 | 8 | 9.25 | <u>7</u> | 10 | 10 | 10 | 10 | 10 | 8 | 8 | 10 | 10 | 10 | 9 | 9 | 9 | 8 | 10 | 1 |
| VOSP position discrimination | 20 | 18 | 18.87 | 19 | 20 | <u>15</u> | 20 | <u>17</u> | 20 | 19 | 20 | 20 | 20 | 20 | 20 | <u>17</u> | 20 | <u>15</u> | 20 | 4 |
| VOSP number location | 10 | 7 | 8.44 | 8 | 10 | <u>5</u> | 8 | 10 | 10 | 10 | 8 | <u>5</u> | 10 | 10 | 10 | <u>4</u> | 9 | 8 | 10 | 3 |
| VOSP cube analysis | 10 | 6 | 8.25 | 8 | 9 | <u>4</u> | 8 | 7 | 9 | 10 | 9 | 10 | 10 | 9 | 10 | <u>5</u> | 9 | <u>5</u> | 10 | 3 |
| *Executive and spatial processing* |  |  |  |  |  |  |  |  |  |  |  |  |  |  |  |  |  |  |  |  |
| TEA: counting without distraction | 7 | 4.2 | 5.53 | <u>2</u> | 5 | 6 | 5 | <u>4</u> | NT | 7 | 5 | 5 | 7 | 5 | 6 | 6 | 7 | 6 | 7 | 2 |
| TEA: counting with distraction | 10 | 2.6 | 3.73 | <u>1</u> | 3 | <u>1</u> | 3 | <u>2</u> | NT | 7 | <u>1</u> | <u>2</u> | 6 | <u>2</u> | 9 | 4 | <u>2</u> | 3 | 10 | 7 |
| Raven's coloured matrices | 36 | 28<sup>†</sup> | 29.00 | 31 | <u>27</u> | 31 | 33 | <u>19</u> | 30 | 34 | <u>24</u> | <u>21</u> | 33 | 34 | 32 | <u>20</u> | 32 | <u>27</u> | 36 | 6 |
| Brixton spatial anticipation | 54 | 28 | 26.94 | <u>21</u> | <u>7</u> | <u>18</u> | 39 | <u>24</u> | <u>23</u> | 31 | 34 | 31 | 30 | 41 | 32 | 30 | <u>21</u> | <u>18</u> | 31 | 7 |
| Trail Making Test A | 24 | 24<sup>†</sup> | 22.88 | <u>19</u> | <u>22</u> | <u>23</u> | 24 | 24 | <u>23</u> | 24 | 24 | 24 | 24 | 24 | 24 | 24 | <u>16</u> | <u>23</u> | 24 | 6 |
| Trail Making Test B | 23 | 17.4<sup>†</sup> | 17.37 | <u>2</u> | 23 | <u>16</u> | 21 | <u>1</u> | <u>5</u> | 23 | 23 | 19 | 22 | 22 | 20 | 19 | 20 | 20 | 22 | 4 |

*Note.* Scores are number of correct responses unless otherwise specified. NT = unavailable for testing; NA = testing was not attempted because patients were non-fluent; TEA = Test of Everyday Attention, elevator counting subtest; VOSP = Visual Object and Space Processing battery. Cut-offs for impairment correspond to two standard deviations below control mean performance (separate from the control sample used in the current study), with impaired scores underlined and in bold. These are taken from control norms from respective tests manuals, unless otherwise specified (see below).
<sup>†</sup> Cut-offs taken from control testing at the \*ANONYMISED\*. Number of controls: Raven's = 20; Trail Making Test = 14.
<sup>‡</sup> Cut-off reflects the 'very slow' classification taken from Kerschensteiner, Poeck and Brunner (1972).
<sup>§</sup> Patients P13 and P14 completed only 4 categories (animals, fruit, birds, breeds of dog) for the test of category fluency, while the other patients completed 8 (animals, fruit, birds, breeds of dog, household objects, tools, vehicles, types of boat).

**Table 2.**
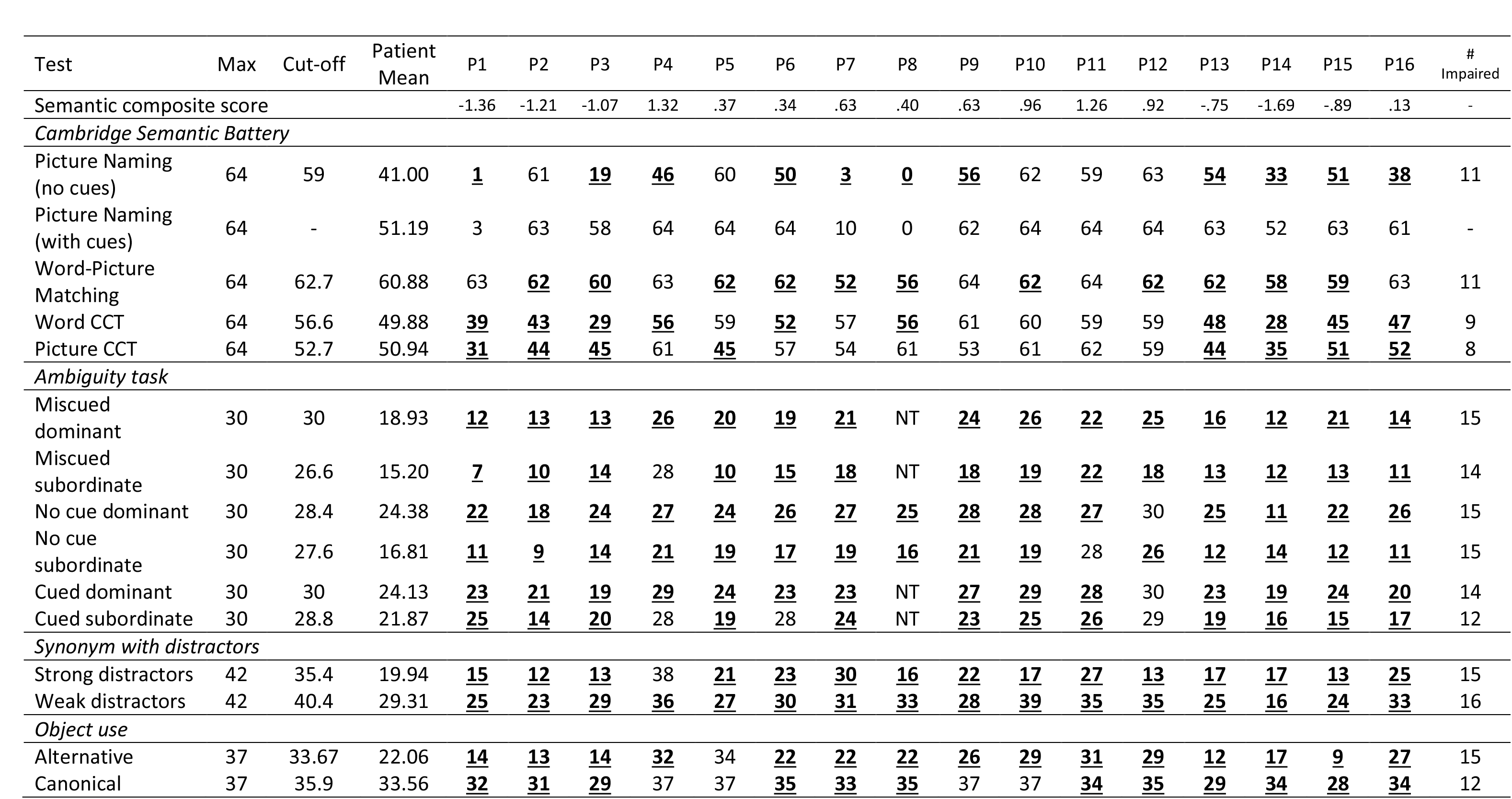
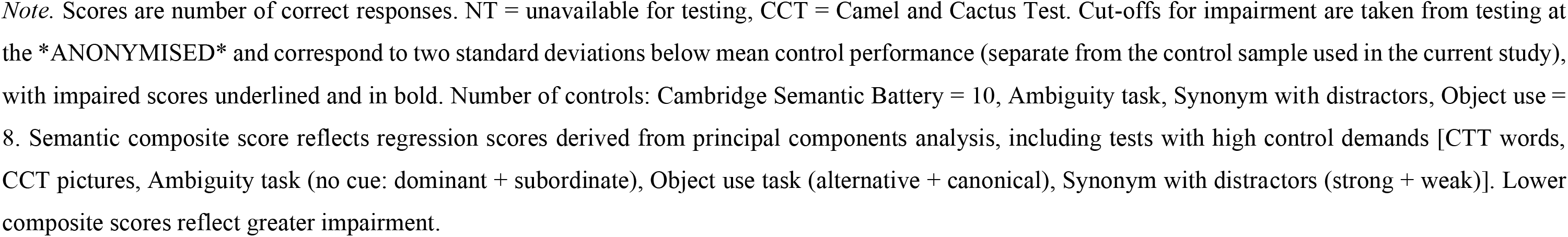
Patient performance on the Cambridge Semantic Battery and tests of semantic control.

| Test | Max | Cut-off | Patient Mean | P1 | P2 | P3 | P4 | P5 | P6 | P7 | P8 | P9 | P10 | P11 | P12 | P13 | P14 | P15 | P16 | # Impaired |
| --- | --- | --- | --- | --- | --- | --- | --- | --- | --- | --- | --- | --- | --- | --- | --- | --- | --- | --- | --- | --- |
| Semantic composite score |  |  |  | -1.36 | -1.21 | -1.07 | 1.32 | .37 | .34 | .63 | .40 | .63 | .96 | 1.26 | .92 | -.75 | -1.69 | -.89 | .13 | - |
| *Cambridge Semantic Battery* |  |  |  |  |  |  |  |  |  |  |  |  |  |  |  |  |  |  |  |  |
| Picture Naming (no cues) | 64 | 59 | 41.00 | <u>1</u> | 61 | <u>19</u> | <u>46</u> | 60 | <u>50</u> | <u>3</u> | <u>0</u> | <u>56</u> | 62 | 59 | 63 | <u>54</u> | <u>33</u> | <u>51</u> | <u>38</u> | 11 |
| Picture Naming (with cues) | 64 | - | 51.19 | 3 | 63 | 58 | 64 | 64 | 64 | 10 | 0 | 62 | 64 | 64 | 64 | 63 | 52 | 63 | 61 | - |
| Word-Picture Matching | 64 | 62.7 | 60.88 | 63 | <u>62</u> | <u>60</u> | 63 | <u>62</u> | <u>62</u> | <u>52</u> | <u>56</u> | 64 | <u>62</u> | 64 | <u>62</u> | <u>62</u> | <u>58</u> | <u>59</u> | 63 | 11 |
| Word CCT | 64 | 56.6 | 49.88 | <u>39</u> | <u>43</u> | <u>29</u> | <u>56</u> | 59 | <u>52</u> | 57 | <u>56</u> | 61 | 60 | 59 | 59 | <u>48</u> | <u>28</u> | <u>45</u> | <u>47</u> | 9 |
| Picture CCT | 64 | 52.7 | 50.94 | <u>31</u> | <u>44</u> | <u>45</u> | 61 | <u>45</u> | 57 | 54 | 61 | 53 | 61 | 62 | 59 | <u>44</u> | <u>35</u> | <u>51</u> | <u>52</u> | 8 |
| *Ambiguity task* |  |  |  |  |  |  |  |  |  |  |  |  |  |  |  |  |  |  |  |  |
| Miscued dominant | 30 | 30 | 18.93 | <u>12</u> | <u>13</u> | <u>13</u> | <u>26</u> | <u>20</u> | <u>19</u> | <u>21</u> | NT | <u>24</u> | <u>26</u> | <u>22</u> | <u>25</u> | <u>16</u> | <u>12</u> | <u>21</u> | <u>14</u> | 15 |
| Miscued subordinate | 30 | 26.6 | 15.20 | <u>7</u> | <u>10</u> | <u>14</u> | 28 | <u>10</u> | <u>15</u> | <u>18</u> | NT | <u>18</u> | <u>19</u> | <u>22</u> | <u>18</u> | <u>13</u> | <u>12</u> | <u>13</u> | <u>11</u> | 14 |
| No cue dominant | 30 | 28.4 | 24.38 | <u>22</u> | <u>18</u> | <u>24</u> | <u>27</u> | <u>24</u> | <u>26</u> | <u>27</u> | <u>25</u> | <u>28</u> | <u>28</u> | <u>27</u> | 30 | <u>25</u> | <u>11</u> | <u>22</u> | <u>26</u> | 15 |
| No cue subordinate | 30 | 27.6 | 16.81 | <u>11</u> | <u>9</u> | <u>14</u> | <u>21</u> | <u>19</u> | <u>17</u> | <u>19</u> | <u>16</u> | <u>21</u> | <u>19</u> | 28 | <u>26</u> | <u>12</u> | <u>14</u> | <u>12</u> | <u>11</u> | 15 |
| Cued dominant | 30 | 30 | 24.13 | <u>23</u> | <u>21</u> | <u>19</u> | <u>29</u> | <u>24</u> | <u>23</u> | <u>23</u> | NT | <u>27</u> | <u>29</u> | <u>28</u> | 30 | <u>23</u> | <u>19</u> | <u>24</u> | <u>20</u> | 14 |
| Cued subordinate | 30 | 28.8 | 21.87 | <u>25</u> | <u>14</u> | <u>20</u> | 28 | <u>19</u> | 28 | <u>24</u> | NT | <u>23</u> | <u>25</u> | <u>26</u> | 29 | <u>19</u> | <u>16</u> | <u>15</u> | <u>17</u> | 12 |
| *Synonym with distractors* |  |  |  |  |  |  |  |  |  |  |  |  |  |  |  |  |  |  |  |  |
| Strong distractors | 42 | 35.4 | 19.94 | <u>15</u> | <u>12</u> | <u>13</u> | 38 | <u>21</u> | <u>23</u> | <u>30</u> | <u>16</u> | <u>22</u> | <u>17</u> | <u>27</u> | <u>13</u> | <u>17</u> | <u>17</u> | <u>13</u> | <u>25</u> | 15 |
| Weak distractors | 42 | 40.4 | 29.31 | <u>25</u> | <u>23</u> | <u>29</u> | <u>36</u> | <u>27</u> | <u>30</u> | <u>31</u> | <u>33</u> | <u>28</u> | <u>39</u> | <u>35</u> | <u>35</u> | <u>25</u> | <u>16</u> | <u>24</u> | <u>33</u> | 16 |
| *Object use* |  |  |  |  |  |  |  |  |  |  |  |  |  |  |  |  |  |  |  |  |
| Alternative | 37 | 33.67 | 22.06 | <u>14</u> | <u>13</u> | <u>14</u> | <u>32</u> | 34 | <u>22</u> | <u>22</u> | <u>22</u> | <u>26</u> | <u>29</u> | <u>31</u> | <u>29</u> | <u>12</u> | <u>17</u> | <u>9</u> | <u>27</u> | 15 |
| Canonical | 37 | 35.9 | 33.56 | <u>32</u> | <u>31</u> | <u>29</u> | 37 | 37 | <u>35</u> | <u>33</u> | <u>35</u> | 37 | 37 | <u>34</u> | <u>35</u> | <u>29</u> | <u>34</u> | <u>28</u> | <u>34</u> | 12 |

*Note.* Scores are number of correct responses. NT = unavailable for testing, CCT = Camel and Cactus Test. Cut-offs for impairment are taken from testing at the \*ANONYMISED\* and correspond to two standard deviations below mean control performance (separate from the control sample used in the current study), with impaired scores underlined and in bold. Number of controls: Cambridge Semantic Battery = 10, Ambiguity task, Synonym with distractors, Object use = 8. Semantic composite score reflects regression scores derived from principal components analysis, including tests with high control demands [CTT words, CCT pictures, Ambiguity task (no cue: dominant + subordinate), Object use task (alternative + canonical), Synonym with distractors (strong + weak)]. Lower composite scores reflect greater impairment.

All patients were impaired on at least one verbal and one non-verbal semantic task. All patients performed close to ceiling level on word-picture matching, reflecting low controls demands. On word and picture versions of the Camel and Cactus Test of semantic association, half of the sample showed impairment. Patients presented with considerable variation in picture naming, although performance was improved by successive phonemic cues in all who were able to name at least one picture. Patients presented with the anticipated impairment in tests manipulating semantic control demands, including difficulty retrieving subordinate conceptual information, susceptibility to cues and miscues, and difficulty rejecting strong thematic distractors.

Principal components analysis of the semantic tasks using oblique rotation revealed two components with Eigenvalues greater than 1 (Table 3). The first component reflected performance on tasks with high semantic control demands: these factor scores were used as a semantic control composite for each participant. Lower scores reflect greater impairment. This semantic factor was positively correlated with performance on the Brixton Spatial Anticipation Test: r_s_(14) = .837, *p* < .001. It did not relate to performance on any other manipulation of executive function (*p* ≥ .200, see supplementary section *Background Neuropsychology*). The second semantic factor loaded on tasks involving object identification.

**Table 3.** Pattern matrix for principal components analysis of semantic aphasia patients’ performance on semantic tests with oblique rotation.

| Task | Component 1 (Eigenvalue = 4.03) | Component 2 (Eigenvalue = 1.52) |
| --- | --- | --- |
| CCT words | **.876** | .083 |
| CCT pictures | **.896** | -.078 |
| Picture naming | .089 | **.877** |
| Word-picture matching | -.062 | **.916** |
| Ambiguity task | **.900** | .057 |
| Synonym judgement task | **.903** | -.154 |
| Object use task | **.801** | .156 |

*Note: Strong loadings for each component are in bold. CCT = Camel and Cactus Test*

## 3. Experiment 1: The effect of cued extrinsic reward on semantic retrieval

### 3.1. Method

#### 3.1.1. Participants

This sample included 16 patients (nine females) with a mean age of 64.4 years (SD = 12.3), a mean age of leaving education of 17.5 years (SD = 2.9), and a mean of 11.3 years (SD = 6.6) since stroke. These patients were compared with 15 controls (12 females) with a mean age of 70.7 (SD = 9.7), and a mean age of leaving education of 18.8 years (SD = 3.9). Patients and controls were matched for age [*U* = 74.0, *p* = .069], and age of leaving education [*U* = 102.0, *p* = .469].

#### 3.1.2. Design

This experiment used a repeated measured design, with all participants making strong and weak thematic associations under the conditions of high and low reward. A three alternative forced choice format was used: participants were asked to select a target word, presented alongside two foils, based on the strongest thematic association to a probe word. The experiment was conducted over two sessions, each consisting of four high and four low reward blocks. Each block contained eight trials split equally across strong and weak associations. High and low reward blocks were alternated. There were 64 trials per session, and 128 trials in total. There was no difference across sessions for accuracy or response time: *p* ≥ .190.

#### 3.1.3. Stimulus properties

Descriptive statistics for Experiment 1 stimulus properties are reported in Supplementary Table 1. Target and probe words were taken from the Edinburgh Associative Thesaurus, a publicly available dataset of associative strength between words (Kiss et al., 1973). Probes and targets were more related in strong than weak association trials: t(67.6) = 42.1, *p* < .001. There were no differences in association strength across blocks [t < 1], or sessions [t < 1].

We examined frequency, imageability and length of the target and probe words. Subtlex-UK (Van Heuven et al., 2014) was used to obtain word frequency. Sources for imageability ratings included the MRC Psycholinguistic Database (Coltheart, 1981), N-Watch (Davis, 2005), The Glasgow Norms (Scott et al., 2019), Bird et al. (2001a; 2001b), Cortese and Fugett (2004a; 2004b), and Davey et al. (2015). Frequency and imageability scores were on 7-point Likert scales, and were averaged when multiple sources were available. Overall, 17% of frequency scores could not be retrieved. Missing scores were largely for compound words such as ‘snooker ball’ or ‘space suit’. Imageability ratings were obtained for all but one word. For both target and probe words, three separate 2×2 ANOVAs were run with frequency, imageability and length as dependent variables, examining effects of reward (high/low) and association strength (strong/weak). These ANOVAs are reported in Supplementary Table 3. All psycholinguistic properties were matched across reward condition and association strength [*p* ≥ .071].

#### 3.1.4. Procedure

Each session was preceded by an instructions phase, which included two practice trials to familiarise participants with the procedure. Stimuli for the main experiment were presented on a laptop using PsychoPy3 (Peirce et al., 2019). Each block was preceded by a graphic, informing the participant that correct answers were each worth 1 point (low reward) or 1,000 points (high reward). Due to impaired reading ability, the researcher read all words aloud to the patients. Patients indicated their response by pointing to the screen, with the researcher pressing the corresponding key. Control participants read the words themselves, and keyed in their own responses. Responses were followed by feedback, informing the participant that they had won either 1 or 1,000 points, or that they were incorrect and had not won any points. If a response was not given within ten seconds, participants were informed that they had not won any points. The prospect of gaining points was abstract and not linked to monetary gain. Each block was followed by self-report ratings of task enjoyment, response confidence, and task focus, each on a 7-point Likert scale. Figure 2 provides a summary of the procedure for Experiment 1.

**Figure 2:**
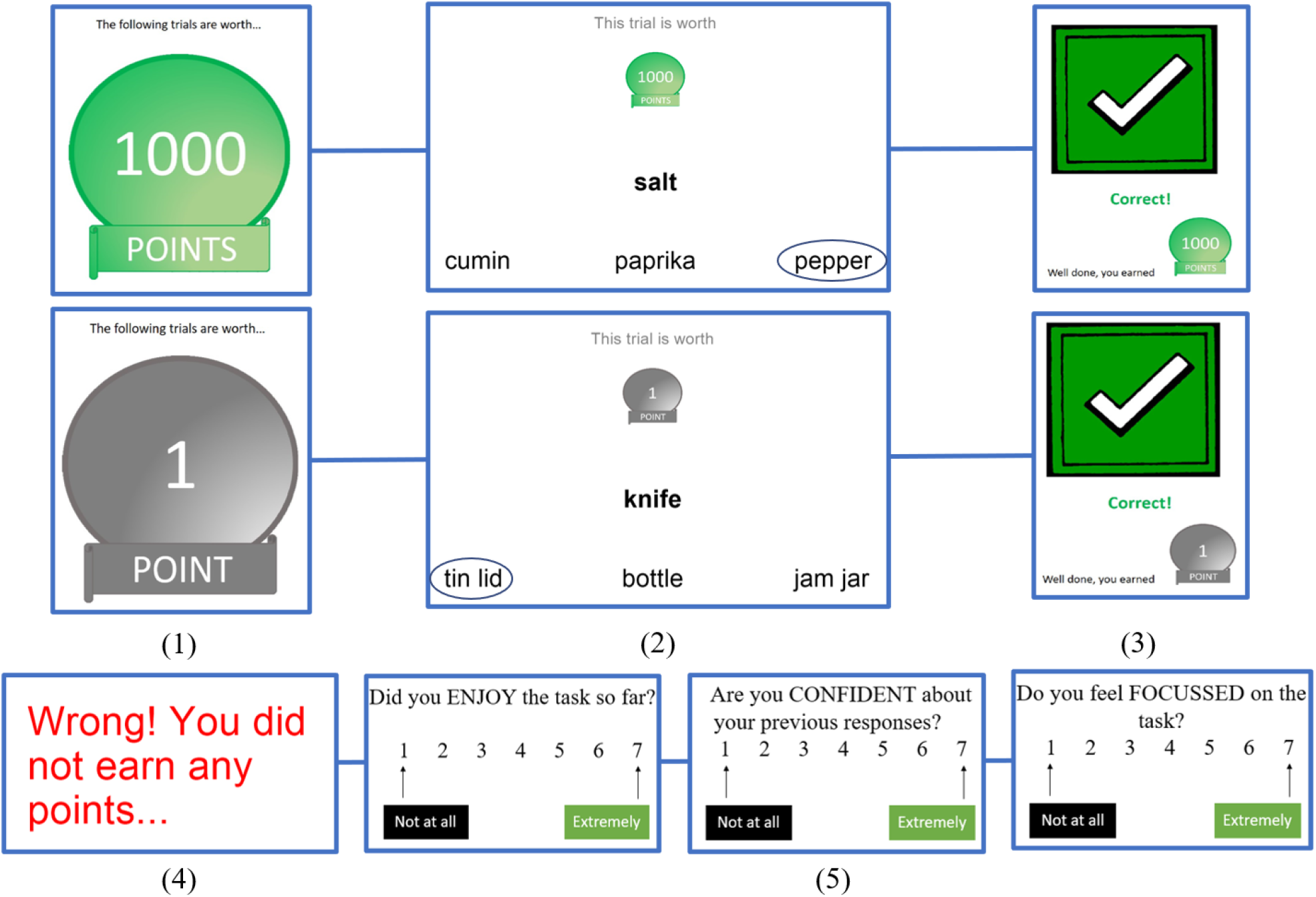
Experiment 1 procedure. (1) Each block was preceded by a high reward or low reward graphic. (2) Participants made thematic associations, either with strong or weak associations. Participants were provided with feedback as to whether their response was (3) correct or (4) incorrect. (5) Following each block, participants completed ratings of enjoyment, confidence, and focus.

#### 3.1.5. Data analysis

Accuracy (proportion of correct responses) was our key dependent measure. As we were specifically interested in effects of reward on patients’ accuracy for weak associations, we first ran a repeated measures ANOVA for the patients alone, observing the effects of reward (high/low) and association strength (strong/weak) as within-subject independent variables. Accuracy was then entered into an omnibus mixed ANOVA, adding group (patients/controls) as a between-subjects variable. Post-hoc contrasts for significant interactions are reported with Bonferroni-correction applied. Mixed ANOVAs were conducted for ratings of enjoyment, confidence, and focus, examining effects of reward and group. Analysis and interpretation of participants’ response time can be seen in Supplementary Table 4.

### 3.2. Results

Figure 3 shows participants’ mean accuracy and self-report ratings across reward condition, group, and association strength. Supplementary Table 5 provides descriptive statistics for Experiment 1.

**Figure 3:**
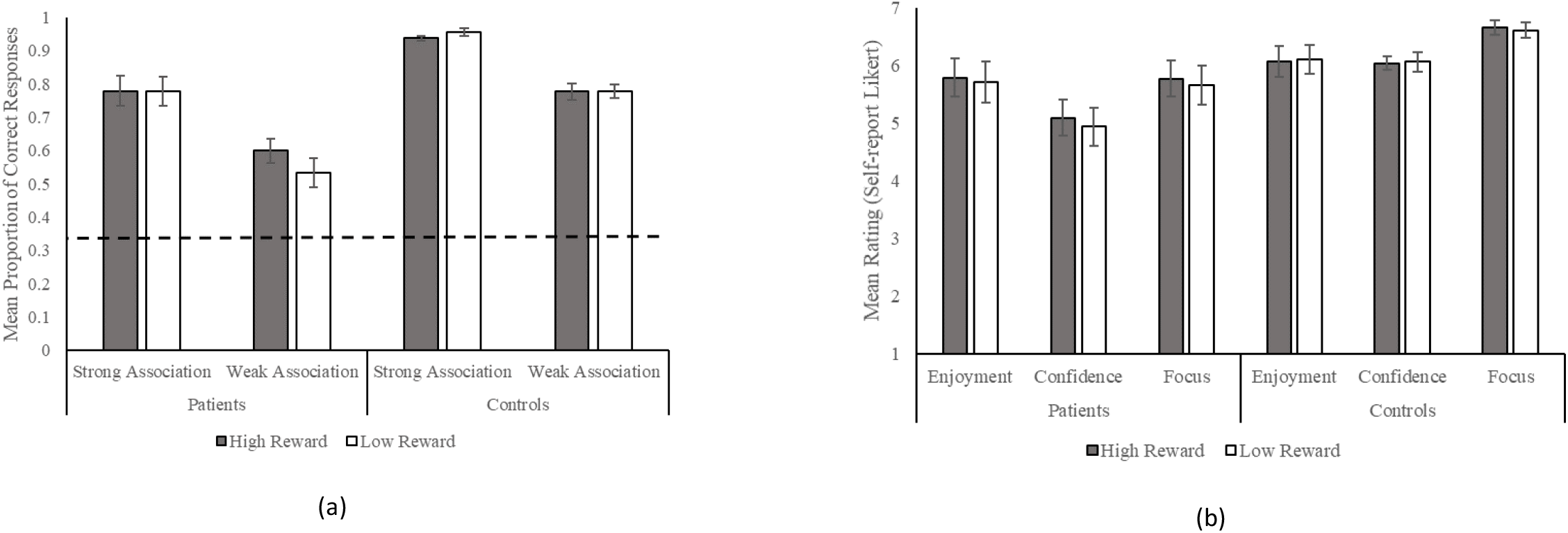
Experiment 1 bar graphs for (a) mean proportion of correct response (dotted line reflects chance performance, .33), and (b) self-report ratings across reward conditions, participant groups, and association strength, with standard error of the mean error bars.

#### 3.2.1. Effects of reward on semantic retrieval in patients

The patient group ANOVA revealed significant main effects of strength [F(1, 15) = 147.7, *p* < .001, η_p_^2^ = .91] and reward [F(1, 15) = 5.4, *p* = .034, η_p_^2^ = .27], and a significant reward by strength interaction [F(1, 15) = 7.0, *p* = .019, η_p_^2^ = .32]. Patients had higher accuracy on high than low reward trials, and on strong than weak association trials. Post-hoc contrasts for the interaction demonstrated that patients were more accurate for weak associations in the high than low reward condition [t(15) = 3.3, corrected *p* = .010]. There was no effect of reward on strong associations [t < 1]^1^. Patients’ semantic control composite scores positively correlated with their overall accuracy [*r*_s_(14) = .70, *p* = .003], reflecting higher accuracy in less impaired patients. There was no association between the semantic composite and the effect of reward [strong: *r*_s_(14) = .16, *p* = .563, weak: *r*_s_(14) = -.08, *p* = .780].

#### 3.2.2. Omnibus ANOVA comparing effects across patients and controls

ANOVA results are shown in Table 4. Controls were more accurate than patients overall. Accuracy was higher for strong than weak association trials. There was a reward by group interaction, with a larger difference in accuracy between high and low reward trials for patients [t(15) = 2.3, corrected *p* = .068] than controls [t < 1]^2^, although neither contrast survived correction. There was a reward by strength interaction, with a greater difference in accuracy between the high and low reward conditions for weak [t(30) = 2.1, corrected *p* = .096] than for strong association trials [t < 1]^3^, although again neither contrast survived correction. The three-way reward by strength by group interaction was not significant.

**Table 4.** Omnibus ANOVA results for all Experiment 1 (extrinsic reward) dependent variables.

| Dependent variable | Main effect/interaction | Results |
| --- | --- | --- |
| Accuracy | Group | **F(1, 29) = 18.9, $p < .001$, $\eta_p^2 = .40^*$** |
| | Reward | $F(1, 29) = 1.3, p = .263, \eta_p^2 = .04$ |
|  | Reward by group | **F(1, 29) = 4.8, $p = .037$, $\eta_p^2 = .14^*$** |
|  | Strength | **F(1, 29) = 215.3, $p < .001$, $\eta_p^2 = .88^*$** |
| | Strength by group | $F(1, 29) = 2.9, p = .100, \eta_p^2 = .09$ |
|  | Reward by strength | **F(1, 29) = 5.6, $p = .025$, $\eta_p^2 = .06^*$** |
| | Reward by strength by group | $F(1, 29) = 2.0, p = .169, \eta_p^2 = .06$ |
| Enjoyment | Group | $F(1, 29) = .6, p = .438, \eta_p^2 = .02$ |
| | Reward | $F(1, 29) = .2, p = .690, \eta_p^2 < .01$ |
| | Reward by group | $F(1, 29) = .6, p = .441, \eta_p^2 = .02$ |
| Confidence | Group | **F(1, 29) = 7.7, $p = .010$, $\eta_p^2 = .21^*$** |
| | Reward | $F(1, 29) = 1.4, p = .244, \eta_p^2 = .05$ |
| | Reward by group | $F(1, 29) = 2.7, p = .112, \eta_p^2 = .09$ |
| Focus | Group | **F(1, 29) = 6.6, $p = .016$, $\eta_p^2 = .19^*$** |
|  | Reward | **F(1, 29) = 5.8, $p = .023$, $\eta_p^2 = .17^*$** |
| | Reward by group | $F(1, 29) = .8, p = .379, \eta_p^2 = .03$ |

*Note: \* reflects a significant result.*

Ratings of enjoyment and confidence were not influenced by reward. Controls reported significantly higher confidence and focus than patients. A main effect of reward was found for focus, with higher ratings in the high than low reward condition. As all ratings were taken at the block level, it was not possible to investigate effects of association strength. There were too few participants in the current sample to assess the relationship between these ratings and accuracy.

#### 3.3. Experiment 1 summary

Experiment 1 studied effects of cued extrinsic reward on semantic aphasia patients’ and controls’ ability to retrieve thematic associations. An ANOVA for the patient group demonstrated that high reward improved accuracy for weak but not strong associations, suggesting that high extrinsic reward can aid the retrieval of semantic associations when semantic control is deficient. Results from the omnibus ANOVA suggest that benefits of extrinsic reward were greater for the patients than for controls, and for weak than strong associations. Self-reported focus was also higher in the high than low reward condition.

## 4. Experiment 2: The effect of self-reference on semantic retrieval

### 4.1. Method

#### 4.1.1. Participants

Experiment 2 included a subset of Experiment 1 participants. This included ten SA patients (six females) with a mean age of 62.4 years (SD = 10.1), a mean age of leaving education of 18.3 years (SD = 3.4), and a mean of 10.1 years (SD = 5.4) since stroke. Eleven control participants (eight females) were included in this sample with a mean age of 69.9 (SD = 10.3), and a mean age of leaving education of 18.9 (SD = 3.6). There was no significant difference between groups for age [*U* = 29.0, *p* = .067], or age leaving education [*U* = 52.0, *p* = .831].

#### 4.1.2. Design

This experiment used a repeated-measures design, with participants making strong and weak thematic associations across ‘self’ and ‘other’ conditions. A three alternative forced choice format was used. Probe pictures were used, as these were viewed as fitting in the context of the shopping game used to reinforce self-referential encoding (explained in *4.1.4.*). Pictures were selected for 28 pairs of semantically related items (e.g., HARP-LUTE, ANT-WASP). For each pair, one picture was allocated to the participant (‘self’) and one to the researcher (‘other’), counterbalanced across participants. Self and other trials were presented in a random order. The experiment was conducted over two sessions, each containing 56 trials. During the first session, participants completed strong and weak associations for one item in each pair. During the second session, participants completed the same associations for the remaining probes. Foils were thematically related to the target and were also kept consistent across both objects in each pair. Session order was counterbalanced. There was no difference across sessions for accuracy or response time: *p* ≥ .259.

#### 4.1.3. Stimulus properties

Descriptive statistics for Experiment 2 stimulus properties are reported in Supplementary Table 2. Association strength between the probe pictures and target words was validated using ratings from an independent sample of healthy adults, on a 7-point Likert scale. Ratings were collected over three surveys, with sample size ranging between 30 and 42. For both probes within each pair, equivalent strong and weak associations were generated. For example, pictures of a HARP and a LUTE were equally strongly and weakly associated with the target words STRINGS and VINYL, respectively. Associations between probes and targets were rated as stronger on strong than on weak association trials: *p* < .001. Association strength within strong and weak association categories was matched across self-reference conditions and sessions: *p* ≥ .793.

Frequency, imageability, and length were examined for the target words (using the same sources detailed in *3.1.3.*). Ratings of frequency and imageability could not be retrieved for 14% and 7% of target words, respectively. This was largely the case for compound words. One-way ANOVAs were run for each factor, looking for effects of association strength (see Supplementary Table 3). No effects of association strength were found [*p* ≥ .195]. Due to counterbalancing, it was not necessary to compare psycholinguistic ratings across conditions or sessions.

#### 4.1.4. Procedure

At the start of both sessions, participants completed the allocation phase ‘shopping game’, intended to reinforce self-referential encoding. Both the participant and researcher had a ‘shopping list’ in front of them, respectively labelled “My shopping list” and “*researcher’s name*’s shopping list”, including pictures and names of each object allocated to them. The researcher and participant took turns finding the items on their lists and placing them into their respective baskets. Participants searched through a pile of laminated pictures, found the next item on their list, and placed it into their basket. Throughout this process the researcher provided verbal prompts to reinforce the allocations (e.g., *“The next item on my list is a bagel, so I’ll put that in my basket. Your next item is a ciabatta, find that one and put it into your basket.”*).

As in Experiment 1, the testing phase was preceded by two practice trials. The testing phase was performed on a laptop using PsychoPy3 (Peirce et al., 2019). The probe picture was presented above the three response options. Participants were asked to identify which of the three words was most thematically related to the probe. Participants indicated their responses in the same way as in Experiment 1 (see *3.1.4.*). Self-reported ratings of response confidence were taken for each trial. Ratings of task enjoyment and focus were not gathered due to the fully randomised design. It was thought that asking participants to self-report enjoyment and focus after each trial may cause frustration and negatively affect enjoyment or focus.

Finally, participants completed an episodic memory test to test for a self-reference recognition memory effect, shown previously in SA (Stampacchia et al., 2019). Participants were presented with 30 images, 10 of which had been allocated to them (“Mine”), 10 which had been allocated to the researcher (“*researcher’s name*”), and 10 which were not present in the allocation or testing phase (“New”). Participants indicated which of these three categories they believed each picture belonged to. The same test was administered after both sessions. A summary of the Experiment 2 procedure can be seen in Figure 4.

**Figure 4:**
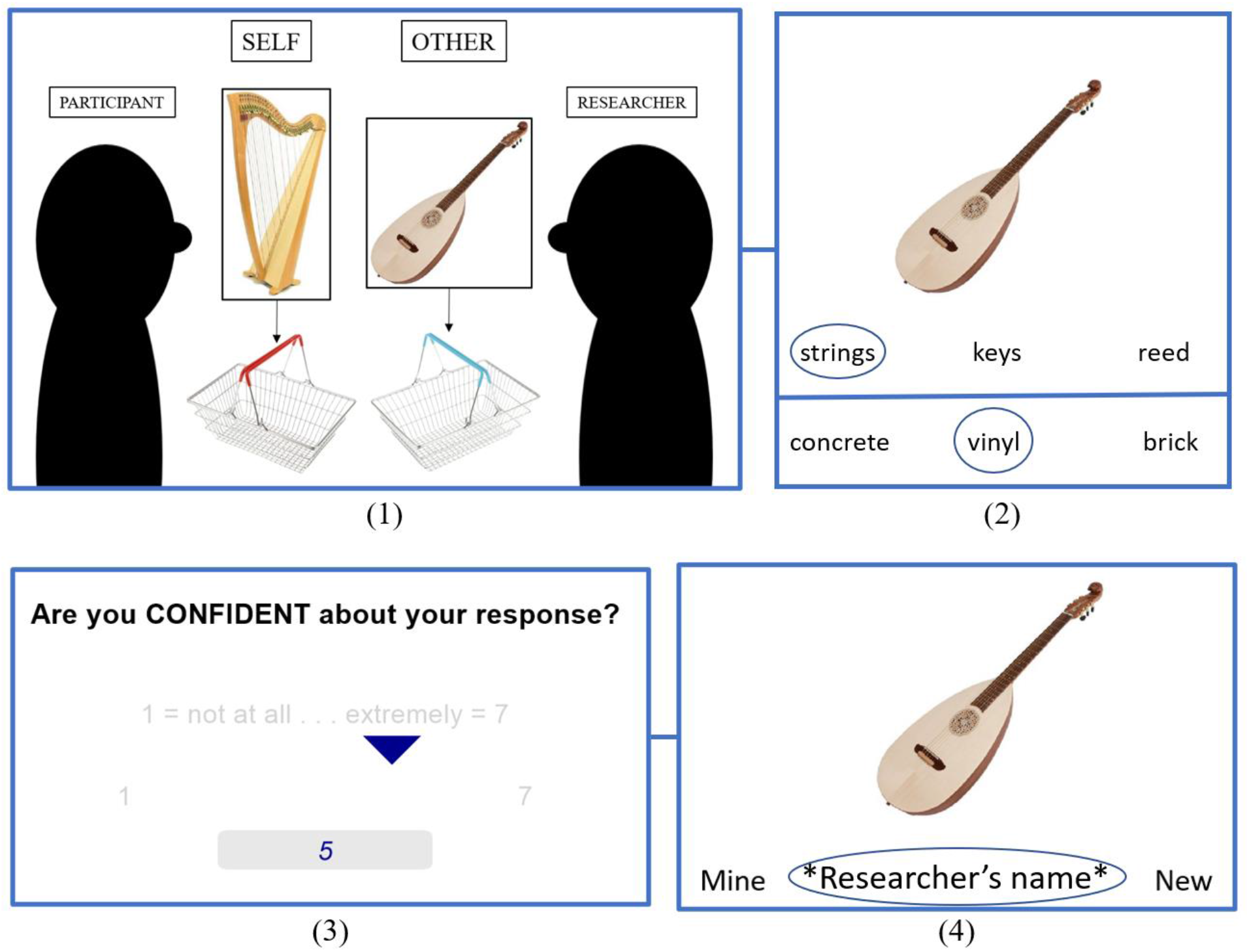
Experiment 2 procedure. (1) One item from each pair of probe pictures was allocated to the participant, and the other to the researcher. (2) Participants completed strong and weak associations for both the self- and other- allocated items in each pair. (3) After each trial participants gave a rating of response confidence. (4) Participants were tested on source memory for 30 pictures.

#### 4.1.5. Data analysis

Accuracy (proportion of correct responses) was our key dependent measure. As benefits of self-reference were only expected for patients on weak association trials, a repeated measures ANOVA was first run for patients only with self-reference condition (self/other) and association strength (strong/weak) as within-subjects variables. Accuracy was then entered into an omnibus mixed ANOVA, adding group (patients/controls) as a between-subjects variable. A mixed ANOVA was conducted for confidence ratings, using the same design as above. Analysis of participants’ response time can be seen in Supplementary Table 4. Results of the episodic memory test were analysed using A’, a non-parametric measure of recognition memory based on the ratio of correct ‘hits’ to false positive responses (Snodgrass & Corwin, 1988). This was calculated for the ‘self’, ‘other’, and ‘new’ conditions. The proportion of correct responses did not vary across sessions for any condition [*p* ≥ .103]. Performance was therefore averaged across both sessions. In cases where memory data was only available for one session (N = 5), data for this session were entered into the analysis. The effects of group and self-reference on A’ were assessed using a mixed ANOVA.

### 4.2. Results

Participants’ A’ scores, mean accuracy, and response confidence across self-reference condition, group, and association strength can be seen in Figure 5. Supplementary Table 6 provides descriptive statistics for Experiment 2. Results for the recognition memory and omnibus ANOVAs are in Table 5.

**Figure 5:**
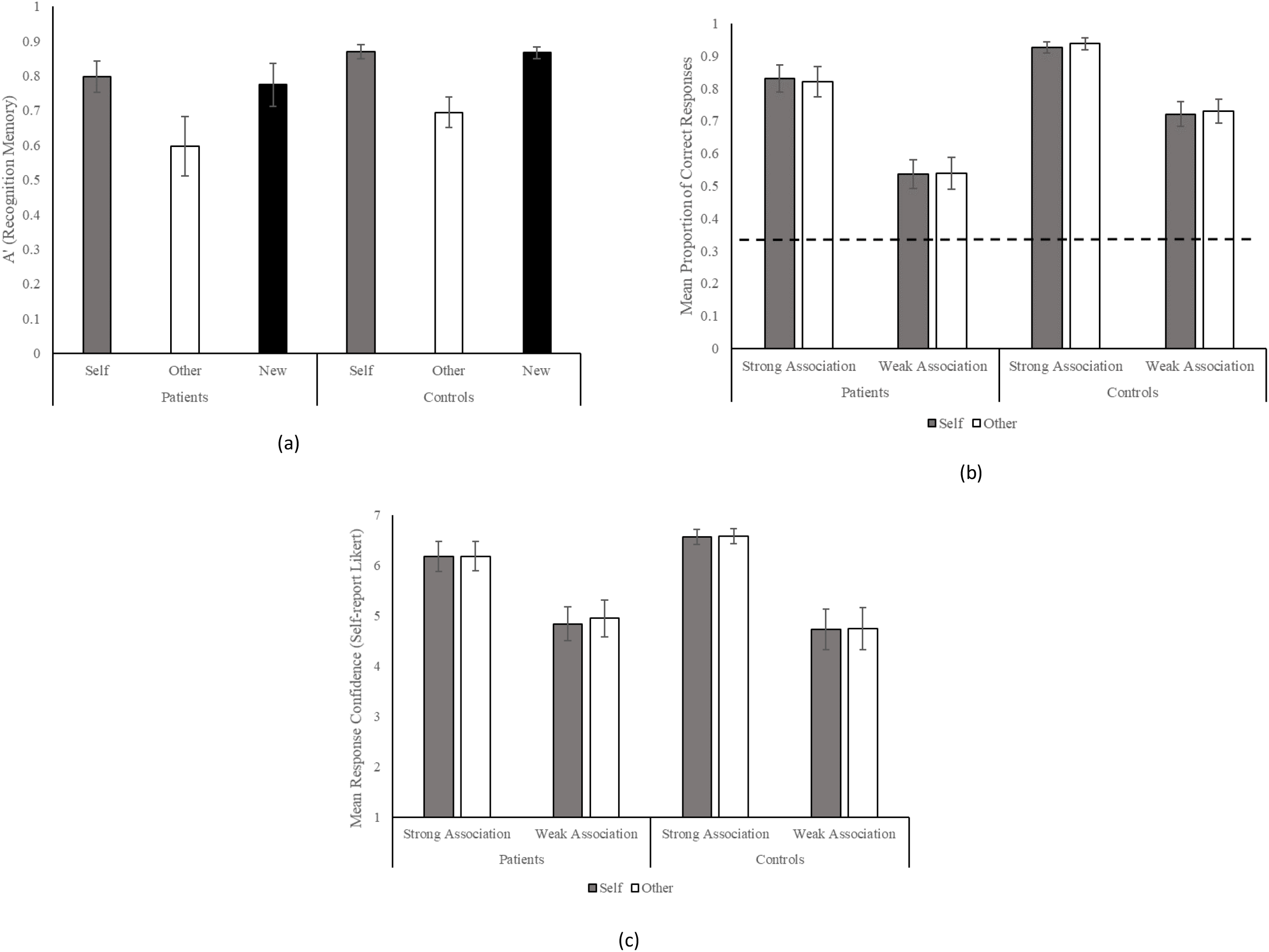
Experiment 2 bar graphs for (a) A’, a non-parametric signal detection measure of recognition memory based on the proportion of correct hits and false positives, (b) mean proportion of correct responses (dotted line = chance), and (c) mean self-report ratings of response confidence, with standard error of the mean error bars.

**Table 5.** Omnibus ANOVA results for all Experiment 2 (self-reference) dependent variables.

| Dependent variable | Main effect/interaction | Results |
| --- | --- | --- |
| A' (recognition memory) | Group | $F(1, 19) = 2.1, p = .168, \eta_p^2 = .10$ |
|  | Self-reference | **$F(1.6, 29.5) = 24.3, p < .001, \eta_p^2 = .56^*$** |
| | Self-reference by group | $F(1.6, 29.5) = .1, p = .861, \eta_p^2 < .01$ |
| Accuracy | Group | **$F(1, 20) = 12.7, p = .002, \eta_p^2 = .38^*$** |
| | Self-reference | $F(1, 20) = .1, p = .767, \eta_p^2 < .01$ |
| | Self-reference by group | $F(1, 20) = .3, p = .595, \eta_p^2 = .01$ |
|  | Strength | **$F(1, 20) = 113.3, p < .001, \eta_p^2 = .87^*$** |
| | Strength by group | $F(1, 20) = 3.7, p = .068, \eta_p^2 = .16$ |
| | Self-reference by strength | $F(1, 20) = .1, p = .803, \eta_p^2 < .01$ |
| | Self-reference by strength by group | $F(1, 20) = .2, p = .677, \eta_p^2 < .01$ |
| Confidence | Group | $F(1, 20) = .1, p = .776, \eta_p^2 < .01$ |
| | Self-reference | $F(1, 20) = .3, p = .589, \eta_p^2 = .02$ |
| | Self-reference by group | $F(1, 20) = .1, p = .746, \eta_p^2 < .01$ |
|  | Strength | **$F(1, 20) = 94.9, p < .001, \eta_p^2 = .83^*$** |
| | Strength by group | $F(1, 20) = 2.9, p = .102, \eta_p^2 = .01$ |
| | Self-reference by strength | $F(1, 20) = .2, p = .688, \eta_p^2 < .01$ |
| | Self-reference by strength by group | $F(1, 20) = .2, p = .675, \eta_p^2 = .01$ |

#### 4.2.1. Effects of self-reference on recognition memory

We report analysis of recognition memory first in order to show the presence of a self-reference memory effect. Planned comparisons of A’ revealed better recognition memory for self-allocated than other-allocated pictures [patients: t(9) = 3.0, *p* = .014, controls: t(10) = 5.6, *p* < .001], and for new than other-allocated pictures [patients: t(9) = 3.3, *p* = .010, controls: t(10) = 4.9, *p* = .001], with no difference between self-allocated and new pictures [patients: t < 1, controls: t < 1] . Both groups therefore showed the expected self-reference memory effect, and effects of novelty.

#### 4.2.2. Effects of self-reference on semantic retrieval in SA patients

For patients’ semantic judgements, ANOVA revealed a significant main effect of association strength [F(1, 9) = 251.1, *p* < .001, η_p_^2^ = .97], with higher accuracy on strong than weak association trials. There was no significant main effect of self-reference [F < 1] or self-reference by strength interaction [F < 1]. Patients’ semantic control composite score positively correlated with overall accuracy [r_s_(8) = .90, *p* < .001], reflecting higher accuracy in less impaired patients.

#### 4.2.3. Omnibus self-reference ANOVAs

In the omnibus ANOVA including both groups (see Table 5), controls were more accurate than patients. There was a main effect of association strength, reflecting higher accuracy for strong than weak association trials. There were no significant effects or interactions involving self-reference. Ratings of confidence were higher for strong than weak association trials.

### 4.3. Experiment 2 summary

Experiment 2 examined the effect of self-reference on SA patients’ and controls’ ability to make thematic associations. As in Experiment 1, controls were more accurate than the patients, and performance was poorer on weak than strong association trials. Greater response confidence was observed for strong versus weak associations. Despite showing a benefit of self-reference for recognition memory, self/other-allocation did not affect the retrieval of thematic associations.

## 5. Discussion

The current study explored the impact of motivation on controlled semantic retrieval in SA patients with multimodal semantic impairment following left frontoparietal stroke. We assessed the impact of performance-contingent extrinsic reward (Experiment 1) and self-referentially encoded pictures (Experiment 2) on patients’ and controls’ ability to retrieve strong and weak thematic associations. As expected, SA patients showed lower accuracy overall. Both groups showed lower accuracy for weak associations, thought to reflect higher semantic control demands. Importantly, extrinsic reward improved SA patients’ but not controls’ accuracy. Self-reference did not impact participants’ semantic performance, despite boosting recognition memory.

SA patients typically show greater semantic impairment for weak associations, when the retrieval of non-dominant information is required (Thompson et al., 2017). In this study, we did not observe the anticipated interactions between group and association strength in accuracy (or response time, see Supplementary Table 4), perhaps because the weak association trials were relatively difficult, eliciting frequent errors even in controls, or because our patient sample included mildly impaired individuals. Future research could address these possibilities by observing effects of parametric manipulations of association strength in SA and/or by including more patients with a wider degree of impairment. Experiment 1 demonstrated improvements in participants’ accuracy for weak but not strong associations following high extrinsic reward. SA patients showed an effect of reward while controls did not, suggesting that when sufficient control over semantic retrieval is harder to achieve, benefits of extrinsic reward are maximised. Anticipation of extrinsic reward may increase preparatory cognitive control, supporting the ability to maintain task-relevant representations and shield against irrelevant information (Goschke & Bolte, 2014). This is consistent with the current finding that high extrinsic reward increased self-reported task focus. Furthermore, explicit knowledge of task goals has been shown to facilitate semantic judgements (Zhang et al., 2021). Reward may benefit semantic control by augmenting goal-maintenance.

Our findings are also consistent with evidence that extrinsic incentives improve performance on domain-general cognitive control tasks (Capa et al., 2013). Neuroimaging research has shown that introducing extrinsic rewards to cognitive control tasks increases activity across MDN regions (Shashidhara et al., 2019), increases functional connectivity between the ventral striatum and MDN (Cubillo et al., 2019), and improves decoding accuracy of MVPA classifiers for task-set information (Etzel et al., 2016). This reflects enhanced coding of task-relevant information, in line with suggestions that extrinsic reward improves goal maintenance (Goschke & Bolte, 2014). The interaction between reward and semantic control seen in the current study may also be attributable to modulation of MDN regions, as well as regions specifically recruited during semantic control. Indeed, MDN regions are recruited during semantic tasks with high control demands (Wang et al., 2020). Future neuroimaging investigations could elucidate the extent to which motivated semantic control is attributable to modulation or recruitment of domain-general versus semantic control regions. Despite their distinct neurobiological underpinnings (Gao et al., 2021), the current findings suggest that modulatory behavioural effects of reward on semantic control mirror those seen for domain-general control.

While there is evidence that semantic and domain-general control are dissociable (Gonzalez Alam et al., 2018), samples of SA patients can show associations between performance on tests of these functions (Thompson et al., 2018). Semantic and executive control substrates are adjacent, such that damage to one system is frequently accompanied by damage to the other (Souter, Wang, et al., 2021; Wang et al., 2018). Accordingly, the current study revealed a positive correlation between semantic ability and performance on the Brixton Spatial Anticipation Test, a complex nonverbal executive test. Patients’ semantic control composite did not correlate with executive measures with verbal requirements including the difference between parts A and B of the Trail Making Test, or with the nonverbal Raven’s coloured progressive matrices. While these null results may reflect a lack of statistical power, our results are sufficient to show that associations between semantic and executive performance are not confined to tests with verbal requirements, consistent with evidence that executive performance is independent of verbal demands in aphasia patients with LIFG lesions (Kendrick et al., 2019; see Chapman et al., 2020 for an alternative view).

The current findings have implications for aphasia rehabilitation. Positive effects of reward are seen in ‘gamification’ strategies to neurorehabilitation (and education more widely), whereby tasks are made more motivating using typical game features, such as rewards and social competition (Landers, 2014). A preliminary investigation demonstrated that gamification may facilitate the rehabilitation of word production following stroke (Romani et al., 2019). The current findings extend this work to show that SA patients can benefit from this strategy, despite deficits of semantic control being accompanied by difficulties in constraining internal representational states in domains beyond semantic cognition, including emotion perception (Souter, Lindquist, & Jefferies, 2021) and episodic memory (Stampacchia et al., 2018). These findings merit further investigation of the use of gamified extrinsic incentives in addressing post-stroke impairments in semantic control. SA patients benefit from external prompts which allude to target concepts, including phonemic cues (Jefferies et al., 2008), context-relevant sentences (Noonan et al., 2010), and emotional cues (Lanzoni et al., 2019). The current findings demonstrate that prompts which do not provide additional information concerning target concepts, such as abstract extrinsic incentives, can confer similar benefits.

The current manipulation of self-reference was intended as a proxy for intrinsic motivation, based on evidence of overlapping behavioural effects of self-reference and reward processing (Sui & Humphreys, 2015a), and overlapping neural substrates underlying self-reference and intrinsic motivation (Tamir & Mitchell, 2012). We found expected effects of self-reference on recognition memory, suggesting that we successfully evoked self-referential encoding, consistent with prior evidence from SA (Stampacchia et al., 2019). Self-reference was not found to modulate semantic retrieval. This null result does not preclude the role of intrinsic motivation in semantic performance; the manipulation in the current study may have been insufficient. In future studies, further tailoring may be required to elicit stronger intrinsic motivators. As intrinsic motivation reflects inherent interest or enjoyment (Mori et al., 2018), it may be beneficial to include stimuli which are specifically of interest to, or belong to, the participant.

### 5.2. Limitations

The current study is limited in so far as we did not measure several constructs related to reward processing. Affective abnormalities including apathy (Fishman et al., 2018) and hypo/hyperarousal (Heilman et al., 1978; Laures et al., 2003) are common following stroke, and could interfere with reward sensitivity. This has been demonstrated in relation to apathy, following damage to subcortical reward processing regions (Rochat et al., 2013). The current study cannot account for these effects. It is worth noting, however, that in the current sample, subcortical and medial regions were relatively intact (see *2.2.*). Future investigations into reward processing in post-stroke aphasia may benefit from measuring apathy, reward sensitivity, and physiological arousal, to better account for effects of these constructs.

### 5.1. Conclusion

The current study demonstrates that extrinsic reward can improve SA patients’ ability to make thematic associations. As with domain-general cognitive control, extrinsic reward may bolster semantic retrieval through increased proactive control. These findings have practical implications for the rehabilitation of post-stroke semantic impairment; language therapy activities for SA patients could be facilitated using a gamification-based approach incorporating external rewards. Effects of self-reference on semantic performance were not observed.

## Data availability statement

Data sharing via a public repository is not possible for the current study, due to insufficient consent. Researchers who wish to access the data should contact the Research Ethics Committee of the York Neuroimaging Centre, or the corresponding author.

## Acknowledgements

We are eternally grateful to the patients, their carers, partners, and families, and the control participants for the time they have given, and continue to give, to support our research. We would also like to thank Dominika Varga for her contribution to stimulus validation, Amelia Shelton, and Marcus Glennon for their contributions to data collection and stimulus validation, Lucilla Lanzoni for her guidance in generating the lesion overlap map used, and Emma Parker, Ellicia Swindells, Annabelle Harding, Chloe Orme, and Cate Correia for their assistance in data collection. Data were provided in part by OASIS (Marcus et al., 2010) for the process of MRI brain extraction.

## Footnotes

1 The assumption of normality was not always met but non-parametric tests elicited the same outcomes. Weak associations: Z = -2.9, corrected *p* = .008; strong associations: Z = -.2, corrected *p* > 1.

2 [Patients: Z = -2.1, corrected *p* = .070, controls: Z = -.4, corrected *p* > 1].

3 [Weak association: Z = -2.4, corrected *p* = .034, strong association: Z = -.7, corrected *p* = .976].

## References

Avants, B. B., Tustison, N. J., Song, G., Cook, P. A., Klein, A., & Gee, J. C. (2011). A reproducible evaluation of ANTs similarity metric performance in brain image registration. NeuroImage, 54(3), 2033–2044. https://doi.org/10.1016/j.neuroimage.2010.09.025.

Bird, H., Franklin, S., & Howard, D. (2001a). Age of acquisition and imageability ratings for a large set of words, including verbs and function words*. Behavior Research Methods, Instruments*, & Computers, 33, 73–79. https://doi.org/10.3758/BF03195349.

Bird, H., Franklin, S., & Howard, D. (2001b). ratings.csv. Retrieved April 4, 2020 from Springer Link: https://link.springer.com/article/10.3758%2FBF03195349#SecESM1.

Botvinick, M., & Braver, T. (2015). Motivation and cognitive control: from behavior to neural mechanism. Annual Review of Psychology, 66, 83–113. https://doi.org/10.1146/annurev-psych-010814-015044.

Bozeat, S., Lambon Ralph, M. A., Patterson, K., Garrard, P., & Hodges, J. R. (2000). Non-verbal semantic impairment in semantic dementia. Neuropsychologia, 9, 1207–1215. https://doi.org/10.1016/s0028-3932(00)00034-8.

Burgess, P. W., & Shallice, T. (1997). The Hayling and Brixton Tests. Bury St Edmunds: Thames Valley Test Company.

Camilleri, J. A., Müller, V. I., Fox, P., Laird, A. R., Hoffstaedter, F., & Kalenscher, T. (2018). Definition and characterization of an extended multiple-demand network. NeuroImage, 165, 138–147. https://doi.org/10.1016/j.neuroimage.2017.10.020.

Capa, R. L., Bouquet, C. A., Dreher, J-C., & Dufor, A. (2013). Long-lasting effects of performance-contingent unconscious and conscious reward incentives during cued task-switching. Cortex, 49, 1943–1954. https://doi.org/10.1016/j.cortex.2012.05.018.

Chapman, C. A., Hasan, O., Schulz, P. E., & Martin, R. C. (2020). Evaluating the distinction between semantic knowledge and semantic access: Evidence from semantic dementia and comprehension-impaired stroke aphasia. Psychonomic Bulletin & Review, 27, 607–639. https://doi.org/10.3758/s13423-019-01706-6.

Coltheart, M. (1981). The MRC psycholinguistic database. Quarterly Journal of Experimental Psychology, 33, 497–505. https://doi.org/10.1080/14640748108400805.

Corbett, F., Jefferies, E., & Lambon Ralph, M. A. (2011). Deregulated semantic cognition follows prefrontal and temporo-parietal damage: evidence from the impact of task constraint on nonverbal object use. Journal of Cognitive Neuroscience, 23(5), 1125–1135. https://doi.org/10.1162/jocn.2010.21539.

Cortese, M. J., & Fugett, A. (2004a). Imageability ratings for 3,000 monosyllabic words. Behavior Research Methods, Instruments, & Computers, 36, 384–387. https://doi.org/10.3758/BF03195585.

Cortese, M. J., & Fugett, A. (2004b). cortese2004norms.csv. Retrieved April 4, 2020 from Springer Link: https://link.springer.com/article/10.3758%2FBF03195585#SecESM1.

Cristofori, I., Salvi, C., Beeman, M., & Grafman, J. (2018). The effects of expected reward on creative problem solving. Cognitive, Affective, & Behavioral Neuroscience, 18, 925–931. https://doi.org/10.3758/s13415-018-0613-5.

Cubillo, A., Makwana, A. B., & Hare, T. A. (2019). Differential modulation of cognitive control networks by monetary reward and punishment. Social Cognitive and Affective Neuroscience, 14(3), 305–317. https://doi.org/10.1093/scan/nsz006.

Davey, J., Cornelissen, P. L., Thompson, H. E., Sonkusare, S., Hallam, G., Smallwood, J., & Jefferies, E. (2015). Automatic and controlled semantic retrieval: TMS reveals distinct contributions of posterior middle temporal gyrus and angular gyrus. The Journal of Neuroscience, 35(46), 15230–15239. https://doi.org/10.1523/JNEUROSCI.4705-14.2015.

Davis, C. J. (2005). N-Watch: A program for deriving neighborhood size and other psycholinguistic statistics. Behavior Research Methods, 37(1), 65–70. https://doi.org/10.3758/BF03206399.

Di Domenico, S. I., & Ryan, R. M. (2017). The emerging neuroscience of intrinsic motivation: A new frontier in self-determination research. Frontiers in Human Neuroscience, 11, 145. https://doi.org/10.3389/fnhum.2017.00145.

Dreisbach, G., & Fischer, R. (2012). The role of affect and reward in the conflict-triggered adjustment of cognitive control. Frontiers in Human Neuroscience, 6(342), 1–5. https://doi.org/10.3389/fnhum.2012.00342.

Enzi, B., de Greck, M., Prösch, U., Tempelmann, C., & Northoff, G. (2009). Is our self nothing but reward? Neuronal overlap and distinction between reward and personal relevance and its relation to human personality. PLoS ONE, 4(12), e8429. https://doi.org/10.1371/journal.pone.0008429.

Etzel, J. A., Cole, M. W., Zacks, J. M., Kay, K. N., & Braver, T. S. (2016). Reward motivation enhances task coding in frontoparietal cortex. Cerebral Cortex, 26(4), 1647–1659. https://doi.org/10.1093/cercor/bhu327.

Fishman, K. N., Ashbaugh, A. R., Lanctôt, K. L., Cayley, M. L., Herrmann, N., Murray, B. J., Sicard, M., Lien, K., Sahlas, D. J., & Swartz, R. H. (2018). Apathy, not depressive symptoms, as a predictor of semantic and phonemic fluency task performance in stroke and transient ischemic attack. Journal of Clinical and Experimental Neuropsychology, 40(5), 449–461. https://doi.org/10.1080/13803395.2017.1371282.

Fröber, K., Pfister, R., & Dreisbach, G. (2019). Increasing reward prospect promotes cognitive flexibility: Direct evidence from voluntary task switching with double registration. Quarterly Journal of Experimental Psychology, 72(8), 1926–1944. https://doi.org/10.1177/1747021818819449.

Gao, Z., Zheng, L., Chiou, R., Gouws, A., Krieger-Redwood, K., Wang, X., Varga, D., Lambon Ralph, M. A., Smallwood, J., & Jefferies, E. (2021). Distinct and common neural coding of semantic and non-semantic control demands. NeuroImage, 236, 118230. https://doi.org/10.1016/j.neuroimage.2021.118230.

Gonzalez Alam, T., Murphy, C., Smallwood, J., & Jefferies, E. (2018). Meaningful inhibition: Exploring the role of meaning and modality in response inhibition. NeuroImage, 181, 108–119. https://doi.org/10.1016/j.neuroimage.2018.06.074.

Goodglass, H., Kaplan, E., & Barresi, B. (2001). The Boston Diagnostic Aphasia Examination. Baltimore: Lippincott, Williams & Wilkins.

Goschke, T., & Bolte, A. (2014). Emotional modulation of control dilemmas: The role of positive affect, reward, and dopamine in cognitive stability and flexibility. Neuropsychologia, 62, 403–423. https://doi.org/10.1016/j.neuropsychologia.2014.07.015.

Hallam, G. P., Whitney, C., Hymers, M., Gouws, A. D., & Jefferies, E. (2016). Charting the effects of TMS with fMRI: Modulation of cortical recruitment within the distributed network supporting semantic control. Neuropsychologia, 93, 40–52. https://doi.org/10.1016/j.neuropsychologia.2016.09.012.

Head, H. (1926). Aphasia and kindred disorders of speech (Vol. II). New York: Cambridge University Press.

Heilman, K. M., Schwartz, H. D., & Watson, R. T. (1978). Hypoarousal in patients with the neglect syndrome and emotional indifference. Neurology, 28(3), 229. https://doi.org/10.1212/WNL.28.3.229.

Hoffman, P., McClelland, J. L., & Lambon Ralph, M. A. (2018). Concepts, control, and context: a connectionist account of normal and disordered semantic cognition. Psychological Review, 125(3), 293–328. https://doi.org/10.1016/j.neuropsychologia.2010.12.034.

Hou, M., Grilli, M. D., & Glisky, E. L. (2019). Self-reference enhances relational memory in young and older adults. Aging, Neuropsychology, and Cognition, 26(1), 105–120. https://doi.org/10.1080/13825585.2017.1409333.

Jackson, R. L. (2021). The neural correlates of semantic control revisited. NeuroImage, 224, 117444. https://doi.org/10.1016/j.neuroimage.2020.117444.

Jefferies, E. (2013). The neural basis of semantic cognition: converging evidence from neuropsychology, neuroimaging and TMS. Cortex, 49, 611–625. https://doi.org/10.1016/j.cortex.2012.10.008.

Jefferies, E., & Lambon Ralph, M. A. (2006). Semantic impairment in stroke aphasia versus semantic dementia: a case-series comparison. Brain, 129, 2132–2147. https://doi.org/10.1093/brain/awl153.

Jefferies, E., Patterson, K., & Lambon Ralph, M. A. (2008). Deficits of knowledge versus executive control in semantic cognition: Insights from cued naming. Neuropsychologia, 46, 649–658. https://doi.org/10.1016/j.neuropsychologia.2007.09.007.

Jefferies, E., Thompson, H., Cornelissen, P., & Smallwood, J. (2019). The neurocognitive basis of knowledge about object identity and events: dissociations reflect opposing effects of semantic coherence and control. Philosophical Transactions of the Royal Society B, 375, 20190300. https://doi.org/10.1098/rstb.2019.0300.

Juechems, K., Balaguer, J., Ruz, M., & Summerfield, C. (2017). Ventromedial prefrontal cortex encodes a latent estimate of cumulative reward. Neuron, 93, 705–714. https://doi.org/10.1016/j.neuron.2016.12.038.

Kay, J., Lesser, R., & Coltheart, M. (1992). Psycholinguistic assessments of language processing in aphasia (PALPA). Hove (UK): Lawrence Erlbaum Associates.

Kendrick, L. T., Robson, H., & Meteyard, L. (2019). Executive control in frontal lesion aphasia: Does verbal load matter? Neuropsychologia, 133, 107178. https://doi.org/10.1016/j.neuropsychologia.2019.107178.

Kerschensteiner, M., Poeck, K., & Brunner, E. (1972). The fluency-non fluency dimension in the classification of aphasic speech. Cortex, 8(2), 233–247. https://doi.org/10.1016/S0010-9452(72)80021-2.

Kiss, G. R., Armstrong, C., Milroy, R., & Piper, J. (1973) An associative thesaurus of English and its computer analysis. In A. J. Aitken, R. W. Bailey, & N. Hamilton-Smith. (Eds.), The Computer and Literary Studies. Edinburgh: University Press.

Kouneiher, F., Charron, S., & Koechlin, E. (2009). Motivation and cognitive control in the human prefrontal cortex. Nature Neuroscience, 12(7), 939–945. https://doi.org/10.1038/nn.2321.

Lambon Ralph, M. A., Jefferies, E., Patterson, K., & Rogers, T. T. (2017). The neural and computational bases of semantic cognition. Nature Reviews Neuroscience, 18(1), 42–55. https://doi.org/10.1038/nrn.2016.150.

Landers, R. N. (2014). Developing a theory of gamified learning: linking serious games and gamification of learning. Simulation & Gaming, 45(6), 752–768. https://doi.org/10.1177/1046878114563660.

Lanzoni, L., Thompson, H., Beintari, D., Berwick, K., Demnitz-King, H., Raspin, H., Taha, M., Stampacchia, S., Smallwood, J., & Jefferies, E. (2019). Emotion and location cues bias conceptual retrieval in people with deficient semantic control. Neuropsychologia, 131, 294–305. https://doi.org/10.1016/j.neuropsychologia.2019.05.030.

Laures, J., Odell, K., & Coe, C. (2003). Arousal and auditory vigilance in individuals with aphasia during a linguistic and nonlinguistic task. Aphasiology, 17(12), 1133–1152. https://doi.org/10.1080/02687030344000436.

Lin, A., Adolphs, R., & Rangel, A. (2012). Social and monetary reward learning engage overlapping neural substrates. SCAN, 7, 274–281. https://doi.org/10.1093/scan/nsr006.

Marcus, D. S., Fotenos, A. F., Csernansky, J. G., Morris, J. C., & Buckner, R. L. (2010). Open access series of imaging studies: Longitudinal MRI data in nondemented and demented older adults. Journal of Cognitive Neuroscience, 22(12), 2677–2684. https://doi.org/10.1162/jocn.2009.21407.

Mekler, E. D., Brühlmann, F., Tuch, A. N., & Opwis, K. (2017). Towards understanding the effects of individual gamification elements on intrinsic motivation and performance. Computers in Human Behavior, 71, 525–534. https://doi.org/10.1016/j.chb.2015.08.048.

Mori, A., Okamoto, Y., Okada, G., Takagaki, K., Takamura, M., Jinnin, R., Ichikawa, N., Yamamura, T., Yokoyama, S., Shiota, S., Yoshino, A., Miyake, Y., Okamoto, Y., Matsumoto, M., Matsumoto, K., & Yamawaki, S. (2018). Effects of behavioural activation on the neural circuit related to intrinsic motivation. BJPsych Open, 4, 317–323. https://doi.org/10.1192/bjo.2018.40.

Noonan, K. A, Jefferies, E., Corbett, F., & Lambon Ralph, M. A. (2010). Elucidating the nature of deregulated semantic cognition in semantic aphasia: Evidence for the roles of prefrontal and temporo-parietal cortices. Journal of Cognitive Neuroscience, 22(7), 1597–1613. https://doi.org/10.1162/jocn.2009.21289.

Noonan, K. A., Jefferies, E., Visser, M., & Lambon Ralph, M. A. (2013). Going beyond inferior prefrontal involvement in semantic control: evidence for the additional contribution of dorsal angular gyrus and posterior middle temporal cortex. Journal of Cognitive Neuroscience, 25(11), 1824–1850. https://doi.org/10.1162/jocn_a_00442.

Notebaert, W., & Braem, S. (2015). Parsing the effects of reward on cognitive control. In T. S. Braver (Ed.), Motivation and Cognitive Control (pp. 105–122). New York, NY: Routledge.

Padmala, S., & Pessoa, L. (2011). Reward reduces conflict by enhancing attentional control and biasing visual cortical processing. Journal of Cognitive Neuroscience, 23(11), 3419–3432. https://doi.org/10.1162/jocn_a_00011.

Parro, C., Dixon, M. L., & Christoff, K. (2018). The neural basis of motivational influences on cognitive control. Human Brain Mapping, 39(12), 5097–5111. https://doi.org/10.1002/hbm.24348.

Peirce, J. W., Gray, J. R., Simpson, S., MacAskill, M. R., Höchenberger, R., Sogo, H., Kastman, E., & Lindeløv, J. (2019). PsychoPy2: experiments in behavior made easy. Behavior Research Methods, 51, 195–203. https://doi.org/10.3758/s13428-018-01193-y.

Raven, J. (1962). Coloured progressive matrices sets A, AB, B. London: H.K. Lewis.

Reitan, R. M. (1958). Validity of the trail making test as an indicator of organic brain damage. Perceptual and Motor Skills, 8, 271–276. https://doi.org/10.2466/pms.1958.8.3.271.

Robertson, I., Ward, T., Ridgeway, V., & Nimmo-Smith, I. (1994). The test of everyday attention. London: Thames Valley Test Company.

Rochat, L., Van der Linden, M., Renaud, O., Epiney, J., Michel, P., Sztajzel, R., Spierer, L., & Annoni, J. (2013). Poor reward sensitivity and apathy after stroke: Implication of basal ganglia. Neurology, 81(19), 1674–1680. https://doi.org/10.1212/01.wnl.0000435290.49598.1d.

Röer, J. P., Bell, R., & Buchner, A. (2013). Self-relevance increases the irrelevant sound effect: Attentional disruption by one’s own name. Journal of Cognitive Psychology, 25(8), 925–931. https://doi.org/10.1080/20445911.2013.828063.

Romani, C., Thomas, L., Olson, A., & Lander, L. (2019). Playing a team game improves word production in poststroke aphasia. Aphasiology, 33(3), 253–288. https://doi.org/10.1080/02687038.2018.1548205.

Samson, D., Connolly, C., & Humphreys, G. W. (2007). When “happy” means “sad”: Neuropsychological evidence for the right prefrontal cortex contribution to executive semantic processing. Neuropsychologia, 45, 896–904. https://doi.org/10.1016/j.neuropsychologia.2006.08.023.

Scott, G. G., Keitel, A., Becirspahic, M., Yao, B., & Sereno, S. C. (2019). The Glasgow Norms: Ratings of 5,500 words on nine scales. Behavior Research Methods, 51, 1258–1270. https://doi.org/10.3758/s13428-018-1099-3.

Shashidhara, S., Mitchell, D. J., Erez, Y., & Duncan, J. (2019). Progressive recruitment of the frontoparietal multiple-demand system with increased task complexity, time pressure, and reward. Journal of Cognitive Neuroscience, 31(11), 1617–1630. https://doi.org/10.1162/jocn_a_01440.

Snodgrass, J. G., & Corwin, J. (1988). Pragmatics of measuring recognition memory: applications to dementia and amnesia. Journal of Experimental Psychology: General, 117(1), 34–50. https://doi.org/10.1037//0096-3445.117.1.34.

Souter, N. E., Lindquist, K. A., & Jefferies, E. (2021). Impaired emotion perception and categorization in semantic aphasia. Neuropsychologia, 162, 108052. https://doi.org/10.1016/j.neuropsychologia.2021.108052.

Souter, N. E., Wang, X., Thompson, H., Krieger-Redwood, K., Halai, A. D., Lambon Ralph, M. A., Thiebaut de Schotten, M., & Jefferies, E. (2021). Mapping lesion, structural disconnection, and functional disconnection to symptoms in semantic aphasia. *bioRxiv*, https://doi.org/10.1101/2021.12.01.470605.

Stampacchia, S., Pegg, S., Hallam, G., Smallwood, J., Lambon Ralph, M. A., Thompson, H., & Jefferies, E. (2019). Control the source: source memory for semantic, spatial and self-related items in patients with LIFG lesions. Cortex, 119, 165–183. https://doi.org/10.1016/j.cortex.2019.04.014.

Stampacchia, S., Thompson, H. E., Ball, E., Nathaniel, U., Hallam, G., Smallwood, J., Lambon Ralph, M. A., & Jefferies, E. (2018). Shared processes resolve competition within and between episodic and semantic memory: evidence from patients with LIFG lesions. Cortex, 108, 127–143. https://doi.org/10.1016/j.cortex.2018.07.007.

Sui, J., He, X., & Humphreys, G. W. (2012). Perceptual effects of social salience: Evidence from self-prioritization effects on perceptual matching. Journal of Experimental Psychology: Human Perception and Performance, 38(5), 1105–1117. https://doi.org/10.1037/a0029792.

Sui, J., & Humphreys, G. W. (2015a). The interaction between self-bias and reward: evidence for common and distinct processes. The Quarterly Journal of Experimental Psychology, 68(10), 1952–1964. https://doi.org/10.1080/17470218.2015.1023207.

Sui, J., & Humphreys, G. W. (2015b). The integrative self: how self-reference integrates perception and memory. Trends in Cognitive Sciences, 19(2), 719–727. https://doi.org/10.1016/j.tics.2015.08.015.

Tamir, D. I., & Mitchell, J. P. (2012). Disclosing information about the self is intrinsically rewarding. PNAS, 109(21), 8038–8043. https://doi.org/10.1073/pnas.1202129109.

Thompson, H. E., Almaghyuli, A., Noonan, K. A., barak, O., Lambon Ralph, M. A., & Jefferies, E. (2018). The contribution of executive control to semantic cognition: Convergent evidence from semantic aphasia and executive dysfunction. Journal of Neuropsychology, 12, 312–340. https://doi.org/10.1111/jnp.12142.

Thompson, H., Davey, J., Hoffman, P., Hallam, G., Kosinski, R., Howkins, S., Wooffindin, E., Gabbitas, R., & Jefferies, E. (2017). Semantic control deficits impair understanding of thematic relationships more than object identity. Neuropsychologia, 104, 113–125. https://doi.org/10.1016/j.neuropsychologia.2017.08.013.

van den Berg, E., Nys, G. M. S., Brands, A. M. A., Ruis, C., van Zandvoort, M. J. E., & Kessels, R. P. C. (2009). The Brixton Spatial Anticipation Test as a test for executive function: Validity in patient groups and norms for older adults. Journal of the International Neuropsychological Society, 15, 695–703. https://doi.org/10.1017/S1355617709990269.

Van Heuven, W. J. B., Mandera, P., Keuleers, E., & Brysbaert, M. (2014). Subtlex-UK: A new and improved word frequency database for British English. Quarterly Journal of Experimental Psychology, 67, 1176–1190. https://doi.org/10.1080/17470218.2013.850521.

Wang, X., Bernhardt, B. C., Karapanagiotidis, T., De Caso, I., Gonzalez Alam, T. R. D. J., Cotter, Z., Smallwood, J., & Jefferies, E. (2018). The structural basis of semantic control: Evidence from individual differences in cortical thickness. NeuroImage, 181, 480–489. https://doi.org/10.1016/j.neuroimage.2018.07.044.

Wang, X., Margulies, D. S., Smallwood, J., & Jefferies, E. (2020). A gradient from long-term memory to novel cognition: Transitions through default mode and executive cortex. NeuroImage, 220, 117074. https://doi.org/10.1016/j.neuroimage.2020.117074.

Warrington, E. K., & James, M. (1991). The Visual Object and Space Battery Perception. Bury St Edmunds: Thames Valley Company.

Wechsler, D. (1997). Wechsler memory scale (3rd ed.). San Antonio, TX: The Psychological Corporation.

Whitney, C., Kirk, M., O’Sullivan, J., Lambon Ralph, M. A., & Jefferies, E. (2011). The neural organization of semantic control: TMS evidence for a distributed network in left inferior frontal and posterior middle temporal gyrus. Cerebral Cortex, 21, 1066–1075. https://doi.org/10.1093/cercor/bhq180.

Yarkoni, T., Poldrack, R., Nichols, T., Van Essen, D. C., & Wager, T. D. (2011). Large-scale automated synthesis of human functional neuroimaging data. Nature Methods, 8(8), 665–670. https://doi.org/10.1038/nmeth.1635.

Yee, D. M., & Braver, T. S. (2018). Interactions of motivation and cognitive control. Current Opinion in Behavioral Sciences, 19, 83–90. https://doi.org/10.1016/j.cobeha.2017.11.009.

Zhang, M., Varga, D., Wang, X., Krieger-Redwood, K., Gouws, A., Smallwood, J., & Jefferies, E. (2021). Knowing what you need to know in advance: The neural processes underpinning flexible semantic retrieval of thematic and taxonomic relations. NeuroImage, 224, 117405. https://doi.org/10.1016/j.neuroimage.2020.117405.

Zhao, X., Jia, L., & Maes, J. H. R. (2018). Effect of achievement motivation on cognitive control adaptations. Journal of Cognitive Psychology, 30(4), 453–465. https://doi.org/10.1080/20445911.2018.1467915.

