## Supplementary Materials for "Motivated semantic control: Exploring the effects of extrinsic reward and self-reference on semantic retrieval in semantic aphasia"

Background Neuropsychology

Patients completed a series of background tests of language, memory, and executive function, the results of which are reported in Table 1 of the main article. While they had common semantic deficits, they were variable in speech fluency and repetition, in line with previous observations (e.g., Stampacchia et al., 2018). Six (of sixteen) patients showed impaired repetition of single words, assessed using a test from the Psycholinguistic Assessments of Language Processing in Aphasia battery (PALPA; Kay et al., 1992). Thirteen and fourteen cases (of fifteen tested) showed impaired forward and backward digit span (Wechsler Memory Scale III; Wechsler, 1997), respectively. Evaluation of spontaneous speech when describing 'The Cookie Theft Picture' (Goodglass et al., 2001) revealed non-fluent speech in nine patients, as consistent with the 'very slow' classification from Kerschensteiner et al. (1972). All patients showed impaired letter fluency (naming words beginning with F, A, and S) and nine (of 13 tested) had impaired category fluency (see Table 1 for categories completed) – three more cases in the group had little speech output and did not attempt these tasks. Visuospatial processing, examined by four subtests of the Visual Object and Space Perception battery (Warrington & James, 1991), was largely preserved. Eleven patients showed impairment on at least one test of executive function, including the elevator counting subtest of the Test of Everyday Attention (Robertson et al., 1994), Raven's Coloured Progressive Matrices (Raven, 1962), The Brixton Spatial Anticipation Test (Burgess & Shallice, 1997), and the Trail Making Test A & B (Reitan, 1958).

Individual patients' performance on the following tests of semantic cognition is reported in Table 2 of the main article. Simple semantic tasks were taken from the Cambridge Semantic Battery (Bozeat et al., 2000). There was considerable variation in performance on Picture Naming, likely due to variable impairment in speech production [Mean (SD) = 64.1% (35.8)]. Though not part of the Cambridge Semantic Battery, successive phonemic cues were provided for words that patients were initially unable to retrieve. With the exception of patients showing near-floor or -ceiling performance, naming improved when phonemic cues were presented [Mean (SD) = 80.0% (36.8), Z = -3.4, $p$ = .001], consistent with the view that SA cases retain knowledge of the names of items that they often fail to retrieve. Word-Picture Matching required patients to match a spoken word to one of ten semantically-related pictures. All patients performed close to ceiling level [Mean (SD) = 95.1% (5.0)], reflecting low control demands. Word and picture versions of the Camel and Cactus Test (CCT) were used to examine the retrieval of thematic associations between concepts. Participants linked a probe word or concept (e.g., ORANGE) to one of four semantically-related options (e.g., target: JUICE, foils: WATER, MILK, WINE). In the word version, the items were presented in print and read aloud by the researcher. There was considerable variation in performance, with half of the sample falling below the normal cut-off for impairment. There was no difference in performance between the word [mean (SD) = 77.9% (16.7)] and picture versions [mean (SD) = 79.6% (14.9)]: Z = -.7, $p$ = .477.

The ambiguity task (Noonan et al., 2010) required patients to make thematic associations between a probe and target word, presented alongside three foils. Each probe was a homonym, with a dominant (e.g., PEN – PENCIL) and subordinate association (e.g., PEN – PIG). These probes were presented alone or following a sentence either cueing the correct interpretation of the probe in relation to the upcoming target (e.g., PEN – PIG: "the labourers cleaned out the pen"), or miscuing the incorrect interpretation (e.g., PEN – PIG: "he signed his name with a fountain pen"). A repeated measures ANOVA assessed the effect of cue condition (no cue, cue, miscue) and dominance (dominant, subordinate). This revealed a significant main effect of cue condition: F(2, 28) = 35.1, $p < .001$, $\eta_p^2$ = .72, and dominance: F(1, 14) = 42.6, $p < .001$, $\eta_p^2$ = .75, and a cue by dominance interaction: F(2, 28) = 8.0, $p$ = .002, $\eta_p^2$ = .37. Planned contrasts confirmed that cues improved performance on subordinate (t(15) = -4.8, $p < .001$) but not dominant trials (t < 1). Conversely, while miscues did not affect performance on subordinate trials (t(15) = 1.6, $p$ = .130), they did impair performance on dominant trials (t(15) = 5.4, $p < .001$)[1].

[1] The assumption of normality was not always met but non-parametric tests elicited the same outcomes. Cue effect for subordinate trials: Z = -3.2, $p$ = .001, cue effect for dominant trials: Z = -.3, $p$ = .752, miscue effect for subordinate trials: Z = -1.7, $p$ = .090, miscue effect for dominant trials: Z = -3.3, $p$ = .001.

The synonym judgement task (Samson et al., 2007) required patients to select which of three possible targets was a synonym of a probe word. Each trial included either a weak distractor, not expected to interfere with retrieval (e.g., probe: HAZARD, target: DANGER, distractor: LIGHT), or a strong distractor, which had a strong thematic association with the probe (e.g., probe: DESERT, target: WILDERNESS, distractor: SAND). The group showed a significant difference in accuracy according to distractor strength: t(15) = -5.2, $p < .001$. In all but one case, accuracy was lower on trials with strong distractors [Mean (SD): 47.5% (17.3)] than with weak distractors [Mean (SD): 69.8% (14.1)].

In the non-verbal object use task (Corbett et al., 2011), patients were presented with a written and spoken action (e.g., "Swat a fly") and pictures of six objects. Patients were required to select which of these objects would be most appropriate for completing the action. In 'canonical' trials, the item typically used (e.g., FLY SWATTER) was present. In 'alternative' trials, the canonical item was not present, but instead an item which could theoretically be used (e.g., MAGAZINE). These alternative trials confer greater control demands by requiring patients to access subordinate conceptual information in relation to the use of target objects, while disregarding irrelevant dominant information (e.g., that magazines are designed to be read). Conversely, targets and probes in canonical trials have strong associations, allowing for relatively automatic retrieval of dominant information. The group showed a significant difference in accuracy between canonical and alternative trials: t(15) = 8.2, $p < .001$. All patients performed better on canonical trials [Mean (SD): 90.7% (8.1)] than on alternative trials [Mean (SD): 59.6% (21.7)].

Overall, every patient showed impairment on at least one verbal and one non-verbal test of semantic cognition, indicating multi-modal impairment in semantic retrieval. Additionally, all patients showed effects of cues, miscues, and strong thematic distractors, as expected for patients with semantic control deficits. This profile is consistent with the classification of SA, and replicate the results of previous studies (e.g., Stampacchia et al., 2018). These impairments occurred in conjunction with variable impairments in language affecting both verbal fluency and repetition. Most cases also showed some evidence of executive dysfunction.

As detailed in section 2.3. of the main article, a ‘semantic control composite score’ was derived using principal components analysis. In order to examine associations between semantic control and executive function, we calculated Spearman correlations between the semantic control composite and each test tapping executive function. This correlation was only found to be significant for the Brixton Spatial Anticipation Test:

- Brixton: **$r_s(14) = .837$, $p < .001$***
- Ravens: $r_s(14) = .338$, $p = .200$
- Test of Everyday Attention (with distraction) – Test of Everyday Attention (without distraction): $r_s(13) = -.275$, $p = .320$
- Trail Making Test B – Trail Making Test A: $r_s(14) = -.024$, $p = .931$
- Digit span backwards – digit span forwards: $r_s(13) = .300$, $p = .278$

The finding that only the Brixton Spatial Anticipation Test was associated with semantic control performance may be partially due to low power in the current sample. Alternatively, this association may reflect the demanding nature of the Brixton test, which requires the identification of rules, updating/switching in accordance with new information, and the inhibition of learned rules. This measure has been shown to be sensitive to executive impairment (van den Berg et al., 2009). Other manipulations used here may be comparatively less demanding. Raven’s Progressive Coloured Matrices requires the discovery of rules, but not the ability to switch between them. The Trail Making Test requires switching, but between well-learned sequences (letters and numbers). Finally, contrasts for the Test of Everyday Attention and digit span tax attentional resources, but present little demand beyond this. Overall, the demands of the Brixton test may be comparable to those of tests implicated in the semantic control composite score, while the other manipulations of executive function used here may be less demanding.

MRI Acquisition

Ten patients (P1 – P10) had MRI scans at *ANONYMISED* using a 3T GE HDx Excite MRI scanner on a T1-weighted 3D fast spoiled gradient echo sequence (TR = 7.8ms, TE = minimum full, flip-angle = $20^{o}$, matrix size = 256 x 256, 176 slices, voxel size = 1.13 x 1.13 x 1mm). All patients were scanned in the chronic stage of stroke (mean (SD) = 8.3 years since stroke (5.4), minimum = 2.5 years). These patients all participated in Experiment 1, while four (P3 – P6) participated in Experiment 2. Structural scans underwent brain extraction in ANTs (version 2.1.0; Avants et al., 2011) using a template from the OASIS Brain Project (https://www.oasis-brains.org/; Marcus et al., 2010). Registration to MNI space was also performed using ANTs. Each patient's lesion location was manually traced in MRIcron.

*Supplementary Table 1. Stimulus properties for Experiment 1.*

| | **Experiment 1 (Reward) – Stimulus Properties [Mean (SD)]** | | | |
|---|---|---|---|---|
| | **High Reward** | | **Low Reward** | |
| | **Strong Association** | **Weak Association** | **Strong Association** | **Weak Association** |
| Probe frequency | 3.72 (.65) | 3.68 (.76) | 3.62 (.64) | 3.75 (.63) |
| Target frequency | 4.08 (.92) | 4.25 (.69) | 4.43 (.59) | 4.26 (.64) |
| Probe imageability | 4.92 (.96) | 5.22 (.95) | 5.05 (.86) | 5.08 (.90) |
| Target imageability | 5.22 (.83) | 5.21 (1.08) | 5.41 (.76) | 5.27 (.91) |
| Probe length | 7.31 (2.07) | 6.41 (2.23) | 7.03 (1.86) | 7.53 (2.53) |
| Target length | 7.22 (2.42) | 6.48 (2.46) | 6.25 (2.26) | 6.03 (2.33) |
| Association strength | 5.70 (.10) | 2.87 (.55) | 5.71 (.10) | 2.98 (.49) |

*Note*: ratings of familiarity, frequency, and imageability taken on a 7-point Likert scale, taken from established databases of linguistic norms. Length corresponds with the numbers of letters in a word. Association strength taken on a 7-point Likert scale, based on ratings taken from the Edinburgh Associative Thesaurus (Kiss et al., 1973).

*Supplementary Table 2. Stimulus properties for Experiment 2.*

| | **Experiment 2 (Self-Reference) – Stimulus Properties [Mean (SD)]** | |
|---|---|---|
| | **Strong Association** | **Weak Association** |
| Target frequency | 4.22 (.73) | 4.05 (.84) |
| Target imageability | 5.27 (.99) | 4.93 (.87) |
| Target length | 6.46 (2.41) | 6.82 (2.83) |
| Set 1 association strength | 5.91 (.64) | 2.78 (.61) |
| Set 2 association strength | 5.89 (.66) | 2.82 (.58) |

*Note*: ratings of familiarity, frequency, and imageability taken on a 7-point Likert scale, taken from established databases of linguistic norms. Length corresponds with the numbers of letters in a word. Association strength taken on a 7-point Likert scale, based on online validation surveys.

*Supplementary Table 3. ANOVAs for Experiment 1 and Experiment 2 psycholinguistic factors.*

| Experiment | Word type | Factor | Main effect/interaction | Results |
|---|---|---|---|---|
| Experiment 1: Reward | Probe | Frequency | Association strength | F(1, 98) = .1, $p$ = .721, $\eta_p^2$ < .01 |
| | | | Reward condition | F(1, 98) < .1, $p$ = .949, $\eta_p^2$ < .01 |
| | | | Reward condition by association strength | F(1, 98) < .1, $p$ = .524, $\eta_p^2$ < .01 |
| | | Imageability | Association strength | F(1, 123) = 1.1, $p$ = .304, $\eta_p^2$ < .01 |
| | | | Reward condition | F(1, 123) < .1, $p$ = .965, $\eta_p^2$ < .01 |
| | | | Reward condition by association strength | F(1, 123) = .7, $p$ = .413, $\eta_p^2$ < .01 |
| | | Length | Association strength | F(1, 124) = .3, $p$ = .600, $\eta_p^2$ < .01 |
| | | | Reward condition | F(1, 124) = 1.2, $p$ = .277, $\eta_p^2$ = .01 |
| | | | Reward condition by association strength | F(1, 124) = 3.3, $p$ = .071, $\eta_p^2$ = .03 |
| | Target | Frequency | Association strength | F(1, 106) < .1, $p$ = .973, $\eta_p^2$ < .01 |
| | | | Reward condition | F(1, 106) = 1.7, $p$ = .194, $\eta_p^2$ = .02 |
| | | | Reward condition by association strength | F(1, 106) = 1.5, $p$ = .224, $\eta_p^2$ = .01 |
| | | Imageability | Association strength | F(1, 124) = .2, $p$ = .624, $\eta_p^2$ < .01 |
| | | | Reward condition | F(1, 124) = .7, $p$ = .416, $\eta_p^2$ < .01 |
| | | | Reward condition by association strength | F(1, 124) = .1, $p$ = .700, $\eta_p^2$ < .01 |
| | | Length | Association strength | F(1, 124) = .3, $p$ = .607, $\eta_p^2$ < .01 |
| | | | Reward condition | F(1, 124) = 2.2, $p$ = .142, $\eta_p^2$ = .02 |
| | | | Reward condition by association strength | F(1, 124) = .4, $p$ = .515, $\eta_p^2$ < .01 |

| | | | | |
|---|---|---|---|---|
| Experiment 2: Self-reference | Target | Frequency | Association strength | F(1, 46) = .6, $p$ = .441, $\eta_p^2$ = .01 |
| | | Imageability | Association strength | F(1, 46) = 1.7, $p$ = .195, $\eta_p^2$ = .03 |
| | | Length | Association strength | F(1, 46) = .3, $p$ = .609, $\eta_p^2$ < .01 |

### Response Time

*Supplementary Table 4. Analysis and interpretation of response time (seconds) in Experiment 1 and Experiment 2*

| Experiment | Main effect/interaction | Results |
|---|---|---|
| Experiment 1: reward | Group | **$F(1, 29) = 80.7$, $p < .001$, $\eta_p^2 = .74$*** |
| | Reward | $F(1, 29) < .1$, $p = .827$, $\eta_p^2 < .01$ |
| | Reward by group | $F(1, 29) = 1.0$, $p = .315$, $\eta_p^2 = .04$ |
| | Strength | **$F(1, 29) = 357.7$, $p < .001$, $\eta_p^2 = .93$*** |
| | Strength by group | $F(1, 29) = 2.7$, $p = .113$, $\eta_p^2 = .08$ |
| | Reward by strength | $F(1, 29) = 3.9$, $p = .057$, $\eta_p^2 = .12$ |
| | Reward by strength by group | $F(1, 29) < .1$, $p = .830$, $\eta_p^2 < .01$ |
| Experiment 2: self-reference | Group | **$F(1, 20) = 18.2$, $p < .001$, $\eta_p^2 = .48$*** |
| | Self-reference | $F(1, 20) = .2$, $p = .662$, $\eta_p^2 = .01$ |
| | Self-reference by group | $F(1, 20) = 1.2$, $p = .278$, $\eta p^2 = .06$ |
| | Strength | **$F(1, 20) = 345.2$, $p < .001$, $\eta_p^2 = .95$*** |
| | Strength by group | $F(1, 20) = 3.2$, $p = .089$, $\eta p^2 = .14$ |
| | Self-reference by strength | $F(1, 20) = .2$, $p = .669$, $\eta p^2 < .01$ |
| | Self-reference by strength by group | $F(1, 20) = .1$, $p = .811$, $\eta p^2 < .01$ |

The observed significant main effects of group on response time reflect that controls responded more quickly than patients in both Experiment 1, and Experiment 2. The observed main effects of strength reflect that all participants responded more slowly for weak than strong association trials in both Experiment 1 and Experiment 2. No effects or interactions of either reward or self-reference for RT were observed. Patients' semantic control composite did not correlate with overall RT in Experiment 1 [$r_s(14) = -.41$, $p = .113$], but did negatively correlate with overall RT in Experiment 2 [$r_s(8) = -.64$, $p = .048$], suggesting slower responses in more impaired patients.

Descriptive Statistics

*Supplementary Table 5. Descriptive statistics for Experiment 1.*

| | **Experiment 1 – Extrinsic Reward** | | | | | | | |
|---|---|---|---|---|---|---|---|---|
| | **Patients (N = 16)** | | | | **Controls (N = 15)** | | | |
| | **Mean** | **SD** | **Minimum** | **Maximum** | **Mean** | **SD** | **Minimum** | **Maximum** |
| *Proportion of correct responses* | | | | | | | | |
| High-strong | .779 | .183 | .28 | .97 | .938 | .031 | .91 | 1 |
| High-weak | .600 | .144 | .31 | .81 | .777 | .095 | .59 | .91 |
| Low-strong | .779 | .176 | .28 | .94 | .956 | .042 | .84 | 1 |
| Low-weak | .533 | .179 | .09 | .81 | .779 | .083 | .66 | .91 |
| *Response time (seconds)* | | | | | | | | |
| High-strong | 5.48 | .931 | 4.51 | 8.03 | 3.02 | .664 | 2.05 | 4.59 |
| High-weak | 6.22 | .913 | 4.96 | 8.01 | 3.94 | .743 | 2.78 | 5.64 |
| Low-strong | 5.33 | .708 | 4.46 | 6.79 | 3.02 | .584 | 2.00 | 4.02 |
| Low-weak | 6.27 | .779 | 5.51 | 7.85 | 4.10 | .631 | 3.01 | 5.28 |
| *Self-reported enjoyment* | | | | | | | | |
| High reward | 5.79 | 1.31 | 2.38 | 7 | 6.08 | 1.03 | 4.00 | 7 |
| Low reward | 5.71 | 1.44 | 2.38 | 7 | 6.10 | .963 | 4.00 | 7 |
| *Self-reported confidence* | | | | | | | | |
| High reward | 5.09 | 1.24 | 2.38 | 7 | 6.04 | .773 | 3.88 | 6.88 |
| Low reward | 4.94 | 1.33 | 2.38 | 7 | 6.07 | .671 | 4.38 | 7 |

| | | | | | | | | |
|---|---|---|---|---|---|---|---|---|
| *Self-reported focus* | | | | | | | | |
| High reward | 5.77 | 1.26 | 2.38 | 7 | 6.67 | .497 | 5.50 | 7 |
| Low reward | 5.66 | 1.34 | 2.38 | 7 | 6.62 | .550 | 5.50 | 7 |

*Note*: The naming of levels of task-based dependent variables is based on the reward condition and association strength of these trials (e.g. High-Strong = high reward, strong association). All self-report ratings were taken on a 7-point Likert scale.

*Supplementary Table 6. Descriptive statistics for Experiment 2.*

| | **Experiment 2 – Self-Reference** | | | | | | | |
|---|---|---|---|---|---|---|---|---|
| | **Patients (N = 10)** | | | | **Controls (N = 11)** | | | |
| | **Mean** | **SD** | **Minimum** | **Maximum** | **Mean** | **SD** | **Minimum** | **Maximum** |
| *Recognition memory – Proportion of hits* | | | | | | | | |
| Self | .810 | .181 | .40 | 1 | .832 | .108 | .60 | .95 |
| Other | .425 | .199 | .15 | .75 | .468 | .169 | .25 | .80 |
| New | .760 | .263 | .10 | 1 | .900 | .087 | .75 | 1 |
| *Recognition memory – Proportion of false positives* | | | | | | | | |
| Self | .365 | .196 | .10 | .75 | .241 | .120 | 0 | .40 |
| Other | .295 | .236 | .10 | .90 | .218 | .108 | .10 | .40 |
| New | .320 | .195 | .05 | .75 | .341 | .130 | .10 | .55 |
| *Recognition memory - A'* | | | | | | | | |
| Self | .797 | .142 | .50 | .94 | .870 | .065 | .73 | .98 |
| Other | .597 | .269 | .09 | .90 | .695 | .148 | .40 | .88 |
| New | .774 | .199 | .36 | .95 | .866 | .057 | .76 | .95 |
| *Proportion of correct responses* | | | | | | | | |
| Self-strong | .831 | .127 | .61 | .96 | .925 | .054 | .79 | .96 |
| Self-weak | .536 | .136 | .32 | .71 | .721 | .120 | .46 | .93 |
| Other-strong | .821 | .139 | .50 | .96 | .938 | .058 | .82 | 1 |
| Other-weak | .539 | .147 | .25 | .75 | .731 | .117 | .50 | .89 |

| | | | | | | | | |
|---|---|---|---|---|---|---|---|---|
| *Response time (seconds)* | | | | | | | | |
| Self-strong | 5.01 | 1.04 | 3.76 | 7.07 | 3.30 | .656 | 2.41 | 4.40 |
| Self-weak | 6.32 | .864 | 4.97 | 7.89 | 4.87 | .812 | 3.55 | 6.12 |
| Other-strong | 4.95 | 1.08 | 3.61 | 7.32 | 3.38 | .629 | 2.48 | 4.49 |
| Other-weak | 6.28 | 1.06 | 4.55 | 7.95 | 5.02 | .686 | 3.91 | 6.24 |
| *Self-reported confidence* | | | | | | | | |
| Self-strong | 6.17 | .993 | 4.25 | 7.00 | 6.56 | .507 | 5.11 | 7.00 |
| Self-weak | 4.84 | 1.07 | 2.75 | 6.15 | 4.73 | 1.33 | 1.50 | 6.07 |
| Other-strong | 6.18 | .943 | 4.11 | 7.00 | 6.57 | .477 | 5.32 | 7.00 |
| Other-weak | 4.95 | 1.22 | 2.96 | 7.00 | 4.74 | 1.40 | 1.61 | 6.14 |

*Note*: A' is a non-parametric signal detection theory statistic, which factors in the proportion of both correct hits and false positive responses. The formula used is from Snodgrass and Corwin (1988). Overall, participants selected 'self' significantly more than 'other' (Z = -3.9, $p < .001$) and 'new' significantly more than 'other' (Z = -3.3, $p = .001$). Given that these responses are believed to reflect genuine recognition memory effects, and given the reported output of A' analysis, this was not considered to be the result of spurious response bias. The naming of levels of task-based dependent variables is based on the self-reference condition and association strength of these trials (e.g. Self-Strong = self-allocated, strong association). Self-report ratings of confidence were taken on a 7-point Likert scale.
